# Reversible superdeformability of hiPSC epithelial cortinoids

**DOI:** 10.1101/2025.09.25.678533

**Authors:** Anirban Jana, Justin Tauber, Adeline Boyreau, Gaëlle Recher, Maxime Feyeux, Basile Gurchenkov, Kevin Alessandri, Pierre Nassoy, L Mahadevan

## Abstract

Epithelial cortinoids, fluid filled shells formed from induced pluripotent stem cells (iPSCs), must accommodate large deformations during growth and morphogenesis. Using inflation–deflation assays and high-resolution imaging, we find that these fluid-filled shells are weakly-pressurized and achieve extreme deformability through reversible soft modes of deformation accommodated by the cytoskeleton. We show that cytoskeletal elements such as actin localized along lateral cell edges undergo tilt and bend instabilities that buffer mechanical load by decoupling apico–basal stretching from lateral extension. These reversible instabilities act as elastic safety valves, permitting large shape changes without loss of epithelial hydraulic and topological integrity. A minimal theoretical and computational model demonstrates how tilt and bend reduce effective resistance to radial thinning and explains the observed pressure–strain softening. Thus, iPSC shells exploit reversible cytoskeletal instabilities as mechanical buffers, enabling robust tolerance of large deformations in developing epithelia.

## 1 Introduction

Human embryo development is a complex multi-cellular process orchestrated by a precise spatial and temporal sequence of self-organization processes including cell growth, division, migration, differentiation events [1]. Embryo development is tightly regulated by a well-defined sequence of shape and morphogenetic changes at the multi-cellular level [2]. At the same time, individual stem cells have been shown to be mechanosensitive: their fate can be influenced by mechanical stress alone or in synergy with biochemical signalling [3, 4]. Direct in-vivo investigation of human embryogenesis remains technically challenging, but advances in stem cell-derived invitro models, along with ethical guidelines, have made organoids powerful tools to dissect the successive stages of early development [5–7].

During implantation of the blastocyst into the maternal uterus, epiblast cells - the progenitors of all embryonic lineages - aggregate into a polarized epithelial shell. These cells establish apico-basal polarity, form tight junctions and generate a central lumen. The nucleation and growth of this lumen is critically dependent on apical actin polarization of specialized structures called apicosomes [8], while subsequent lumen expansion occurs osmotically [9]. To recapitulate these events in-vitro, numerous approaches have been developed using either embryonic (ES) or human induced pluripotent stem cells (hiPSC) [7, 9–11]. These methods promote the self-assembly of closed spherical pseudo-stratified epithelia in carefully controlled mechano-chemical environments [12]. For this process to be reproducible, although adhesion to an extra-cellular matrix environment is not a requirement, the rheological properties or the matrix are essential, as they determine whether iPSC organize into lumen-containing shells instead of solid spheroids [13, 14].

Following implantation, the first major morphological transition is the conversion of an epiblast to an amnion [15]. From a physical viewpoint, it remains unclear how a spherical pseudo-stratified epithelium transforms into a flaccid asymmetric amniotic sac composed of squamous amniotic ectoderm on one side and columnar epiblast at the other. It has been suggested that the trophectoderm-derived tensile forces drive this transformation by reshaping the epiblast into a cup-like shape [12]. However, the mechanical properties of *in vitro* epiblast-like models have not yet been well characterized.

In this work, we devise a quantitative customized nano-inflater to probe the mechanics of iPSC-derived epithelial cortinoids. We show that these shells are weakly pressurized during growth, in contrast with other types of epithelial shells, and they exhibit ultra-low elasticity and super-deformability - unveiling a previously unrecognized deformation mechanism in epithelia.

## 2 Results

### 2.1 Morphological and dynamical features of compartmentalized hiPSC cortinoids

We encapsulated hiPSCs in liquid core capsules made of a nutrient-permeable hydrogel envelope using a microfluidic co-extrusion technique previously developed by us and others [11, 16–18] (Fig.1a). Our routine protocol, which relies on the Rayleigh-Plateau instability, produces capsules with a diameter of 250 ± 50 µm. This compartmentalization enables facile and harmless manipulation of individual capsules. To provide a niche-like environment, Matrigel, an ECM mixture, is co-injected with the cell suspension at a volume fraction required to form a continuous matrix layer anchored to the inner wall of the capsule [19]. Following encapsulation at low seeding density (∼ 5-10 cells/caps), we observed that hiPSCs readily self-assemble into a cortinoid, i.e, a closed spherical epithelial shell, around a central lumen. We monitored the dynamics over days using bright field imaging techniques [20, 21] (Fig.1b and Supp. Video 1). From image analysis (see methods), we derive the shell radius and thickness as a function of time (Fig.1c). Surprisingly, both parameters increase monotonically, as observed in [22] and in contrast to lumenized assemblies of most other epithelial cell types (MDCK, Caco2, and MCF10A) for which the increment in lumen radius is concomitant with thinning of the cell layer [23–26]. Following luminogenesis, an epithelial apico-basal polarity is acquired as early as day 2, as marked by the tight junction protein ZO-1 on the apical side. The cortinoids conserve a columnar epithelium conformation till day 4-5. This epithelium morphology is stable and evolves slowly toward a pseudostratified epithelium [27] on day 6 (Fig.1d), as expected for epiblasts [28]. Although the thickness of the shell reaches almost 50 um at day 6, this value is compatible with the architecture of a pseudo-stratified epithelium [29]. It also qualitatively suggests that this columnar hiPSC shell is not strongly pressurized, contrary to cuboidal MDCK or squamous MCF-DCIs.com shells, where osmotic pressure imbalance due to ion secretion by cells and subsequent passive transport of water into the cavity lead to the establishment of hydrostatic lumen pressure [30–35].

### 2.2 hiPSCs cortinoids are weakly pressurized

The hiPSC cortinoids are encapsulated in a protective hydrogel hollow sphere, which serves us to stick them on a substrate coated with poly-lysine. Using a glass micropipette (diameter ∼ 2-2.5 µm) handled with a micromanipulator under a microscope, the alginate capsule and the shell inside are poked (Fig.1 e). The pipette creates a circular channel of size *d*_*pip*_ across the cell layer, between the inside and the outside (Fig.1 f). This allows us to probe the luminal pressure *P*_*L*_ − *P*_0_ using the Hagen-Poiseuille relationship [35]: Δ*P* = ℛ_*H*_ *Q* where ℛ_*H*_ is the hydraulic resistance of the fluid circuit, and *Q* is the flow rate across the pipette. ℛ_*H*_ is strongly dependent on the profile of the pipette at the tip, and on the viscosity of the intra-luminal liquid *η*, which is taken to be equal to that of water ∼ 10^3^*Pa* − *s*. ℛ_*H*_ is derived by accurately measuring the pipette profile close to the tip (Supp fig. A1).We monitored the lumen volume variation for different initial lumen volumes ranging from 0.05 to 0.35 nL (Fig.1 g and Supp. Vid 2-3). With ℛ_*H*_ =8 × 10^16^ Pa-s-m^−3^, we found that the hydrostatic luminal pressure is always lower than 15 Pa, with an average value around 5 Pa, irrespective of the size and age of the shells (Fig.1 h-i). This estimate is an upper value of the intralumen pressure by assuming that the contact between the cell layer and the pipette is leak-free. We further checked that, upon retrieval of the pipette, lumen volume outflow stopped almost instantaneously, indicating a fast-healing capacity (Extended fig. 1 and Supp. vid 2). Altogether, these experiments demonstrate that our hiPSC cortinoids are very weakly pressurized, suggesting a low tension in the epithelium. From this point of view, the system exhibits an unusual property, since shells or acini are commonly considered as pressurized after luminogenesis or above a critical cavity size [35] [9].

**Fig. 1.**
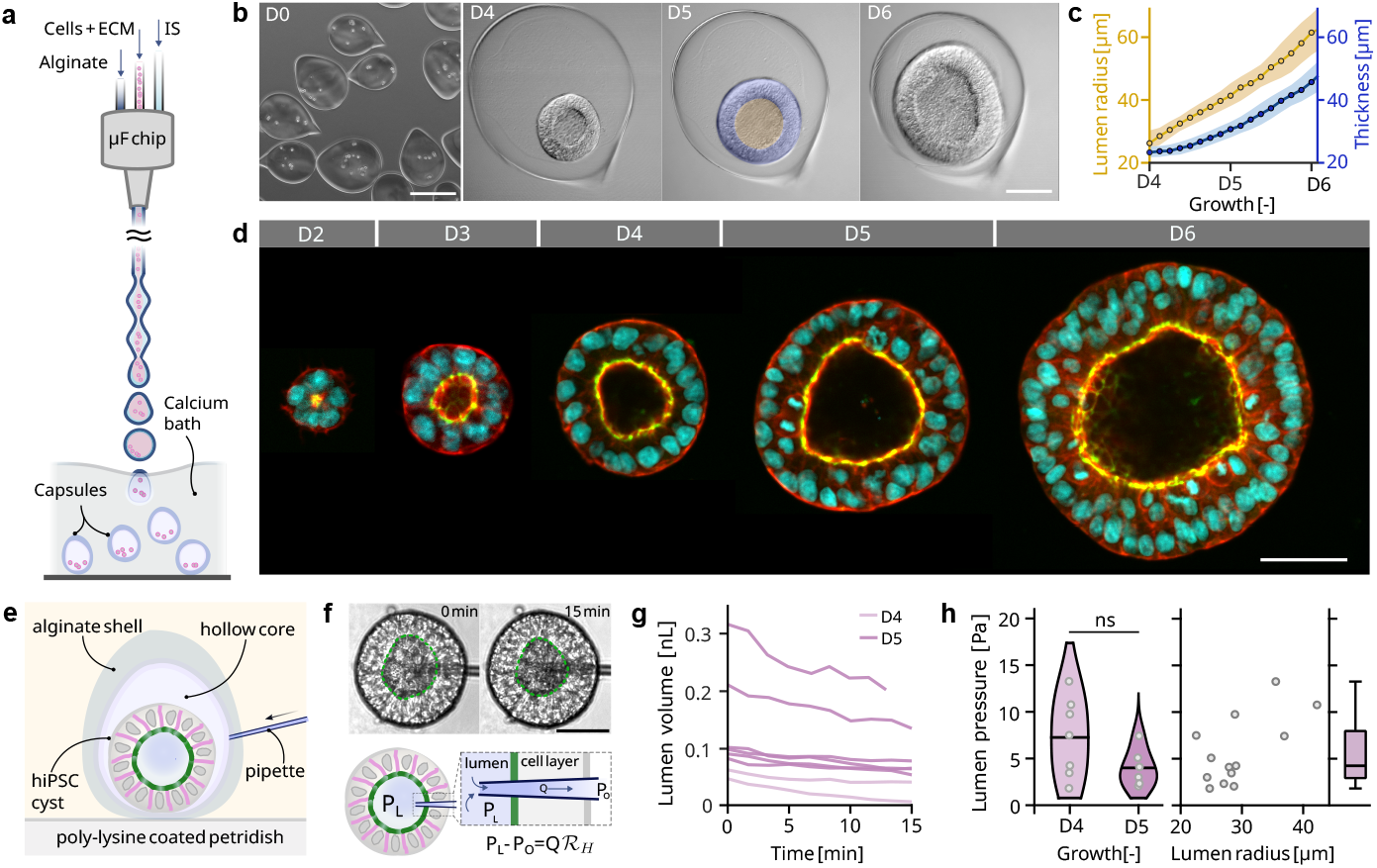
Morphological and dynamical properties of iPSC shells. [a] Schematic of the cellular capsule technology. ECM:Extra-cellular matrix, IS: Intermediate solution, AL:solution of alginate. [b] Bright-field image of capsules taken on day D 0 (day of encapsulation - Scale bar=200 µm), D 4, D 5, D 6 (Scale bar = 50µm). [c] Lumen radius and thickness as a function of time. n=5. [d] Confocal micrographs of hiPSC shells cultured in Matrigel laden alginate capsules from day 2 to day 6. Nuclei in cyan (DAPI 405), ZO-1 tight junctions in green (EGFP 488), cytoskeletal actin in red (Phalloidin 568). Scale bar=50 µm. [e] Schematic of a pipette inserted into a hiPSC cortinoid inside a capsule immobilised on a poly-lysine coated dish. [f] Top: Snapshots of a cortinoid with a pipette inserted in the lumen for 15 minutes. Bottom: Schematic of the lumen fluid drainage through the pipette. *P*_*L*_ and *P*_*O*_ are the pressure inside the lumen and at the free end of the pipette (outside) respectively. [g] Time evolution of the lumen volume for assemblies of different sizes and at different stages of growth after insertion of a pipette. [h] Left: Lumen pressure measured at different stages of growth for cortinoids of different sizes. Violin kernel bandwidth = 0.2. Solid line in each violin plot shows the mean value. p=0.1143 from standard Independent t-test. ns= not significant. For D4, n=7 and D5, n=6; Middle: Lumen pressure as a function of initial lumen radius. Spearman’s correlation coefficient = 0.516. Right: Box plot shows the median (dark line), 25th and 75th quartile (the box), and maximum and minimum (whiskers).

### 2.3 Upon inflation, hiPSC shells remain impermeable, exhibit an elastic behaviour and are super-deformable

We then evaluated the stress response of the cortinoid by injecting fluid into the lumen, leading to a controlled inflation. This approach is the micro-scale analogue of an approach originally devised to investigate the mechanical properties of animal viscera [36]. We first injected an osmotically balanced fluid with a fluorescent dye inside the luminal cavity to test for permeability or leaks (Fig.2 a and Supp. video 4). No leakage is observed at the insertion point of the pipette in the cell layer, confirming fast sealing of the epithelium. More importantly, from the radial profile of fluorescence monitored over about 5 min from the centre to the outer edge of the shell, we were unable to detect any increase in fluorescence outside the shell, indicating that the epithelium retains its integrity and its role as an impermeable barrier (Fig.2 b). Due to incompressibility of water in the pressure range of interest, the deformation of the shell is thus directly related to the volume of fluid injected. In our case, the initial lumen volume of the shells is of the order of 0.2 nL. Since standard flow-meters have a range and resolution that are insufficient to measure flows as low as a fraction of nL/min, the flow rate is derived by monitoring the time evolution of the lumen radius *r* when a pressure *P*_*A*_ (of the order of kPa) is applied. Once the pipette is inserted, the lumen acts as a closed non-pressurized system (*P*_*L*_ ≈ 0). The pressure difference *P*_*A*_ − *P*_*L*_ drives a liquid flow into the lumen. The rate of increase in lumen volume is 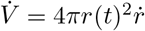, where *r* is the lumen radius and *P*_*L*_ builds up as:

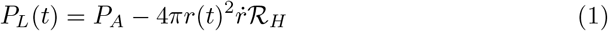

where ℛ_*H*_ is the hydrodynamic resistance of the system (see previous section and Supplementary sec. A1). Due to the resistance of the inflating epithelium, pressure is expected to build up in the lumen and the flow rate is expected to decrease until *P*_*L*_ = *P*_*A*_, leading to a steady state. Conversely, in the inflated state, if *P*_*A*_ is reset to 0, the pressure difference is reversed, leading to an outflow of liquid and a deflation of the cortinoid.

**Fig. 2.**
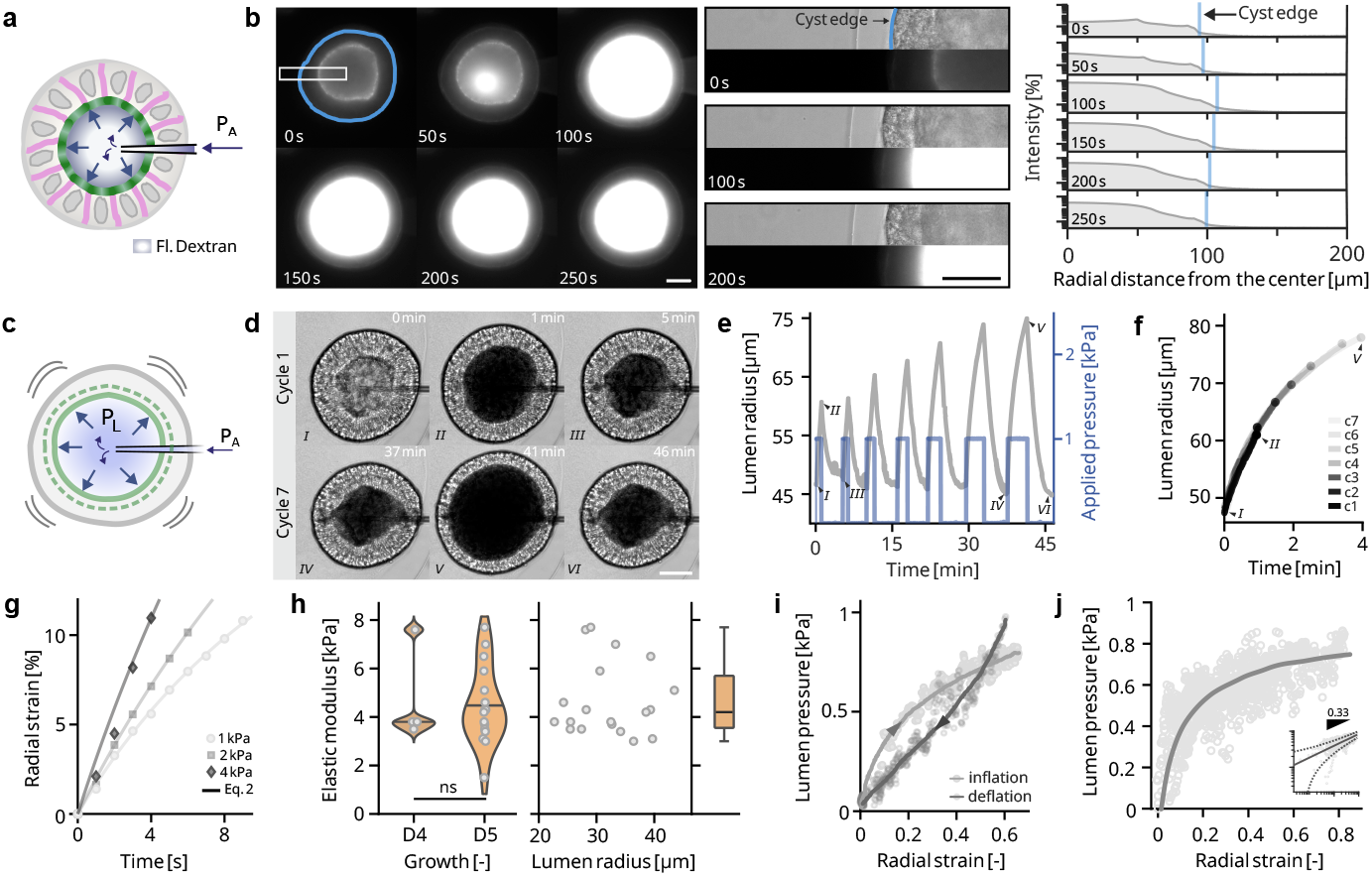
Impermeability and mechanical properties of cortinoids upon inflation. [a] Schematic representation of the experiment of the inflation of a hiPSC shell with fluorescent dextran [b] Left: Grayscale fluorescence snapshots (with FITC-Dextran 20 kDa at 0.1 % as a green fluorescent dye) during inflation. At t=0 s, the fluorescent ring around the lumen is the fluorescent ZO-1 tight junction. Scale bar=50 µm; Middle: Bright-field and Fluorescence micrographs showing simultaneously the cortinoid edge and lumen to quantitatively characterize permeability properties. Scale bar=50 µm; Intensity profiles across the cell layer as a function of radial distance from the center in the region shown in the white box outlined in the image. The shell edge is marked by the blue line. [c] Schematic indicating the notations in a typical inflation experiment where the radial strain is followed with time. [d] Bright-field snapshots of a cortinoid that inflates and relaxes over multiple cycles at a constant pressure step of 1 kPa. From first cycle, the lumen becomes darker due to black dye added to the inflation fluid to increase optical contrast (see Methods for details). Scale bar=50 µm [e] Lumen radius (left Y axis) as a function of time when multiple steps of constant pressures (right Y axis) are applied for various durations. Numbers (in Roman numerals) highlight the time-points corresponding to snapshots shown in [d]. [f] Time evolution of lumen radius for the 7 inflation cycles (c1-c7) shown in [e]. [g] Temporal evolution of luminal radial strain in the low strain regime (*<*10 %) for different constant pressure steps. Experimental data are shown as filled shapes. Numerical solutions of equation (2) (see text for details) at the different applied pressures *P*_*A*_ are shown as solid lines.[h] Left: Elastic modulus of cortinoids on day 4 (n=4) and day 5 (n=15 derived from the curves corresponding to strain *<* 10 %. Markers denote raw data and solid lines in the violins show the mean for each case. Violin kernel bandwidth = 0.2. p-value *>* 0.1 from Mann-Whitney test. ns=non significant. Right: Elastic modulus as a function of the initial lumen radius of the cortinoids (n=19). The box plot shows the median (black line), 25th and 75th quartile (the box), and maximum and minimum (whiskers). [i] Variation of the lumen pressure with the luminal radial strain over a cycle of inflation and deflation (cycle 7 shown in [d]). Solid lines are visual guides. [j] Lumen pressure as a function of radial strain (n=4 cortinoids and n=10 inflations). Solid line is a visual guide. Snippet shows the variation in log scale with a 0.33 power fit.

We devise an experimental procedure in which a constant pressure step (*P*_*A*_) is applied and maintained for an increased duration at each cycle. Variations of the lumen radius are monitored (fig.2 d-e). First, whatever the duration of application of *P*_*A*_, we observe that the lumen relaxes to its initial size upon release of pressure and deflation, indicating reversibility of the deformation (Supp. video 5). However, the deformation reached at the end of the pressure step may vary from about 25% for 1 minute duration (cycle 1 - fig.2 d-e) to 80% when the pressure step is sustained for around 4 minutes (cycle 7 - fig.2 e). By superimposing the different traces of radius with time, there is complete reproducibility but the absence of any steady state (fig.2 f). To extract mechanical properties, we first focus on the low-deformation regime (radial strain *<* 10%). By modelling the system as a thick elastic shell of Young’s modulus *E* undergoing inflation at a constant applied pressure, we derive a constitutive equation (details in Supp. fig A1)

**Fig. 3.**
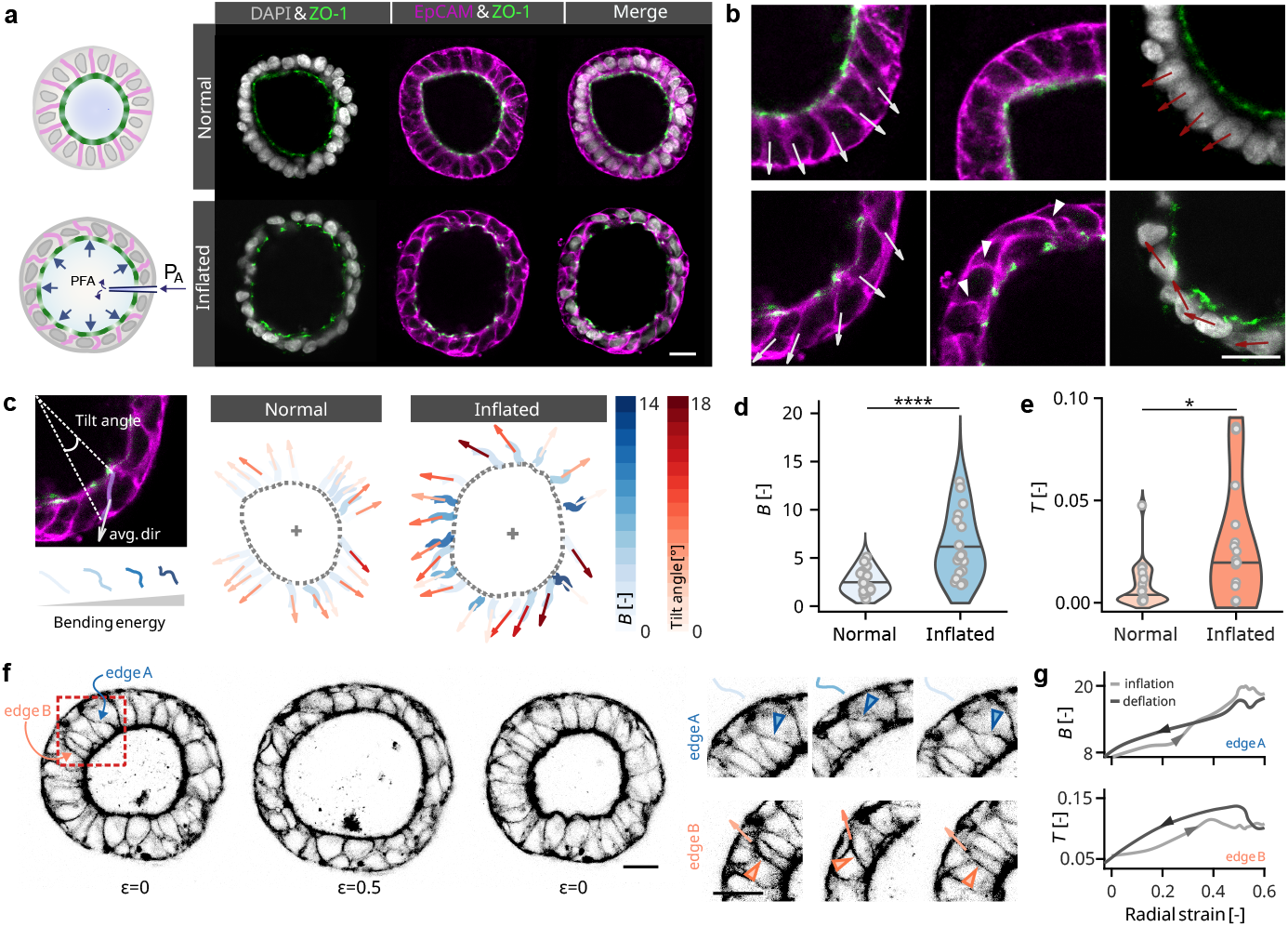
Cell edges bend and tilt at large deformations. [a] Schematic of the protocol to fix a cortinoid in an inflated state with paraformaldehyde (PFA) and corresponding confocal images of the equatorial plane of a cortinoid in the normal resting state (top row) and in the inflated state (bottom row). Nuclei in grayscale (DAPI), ZO-1 tight junctions in green (GFP) and basolateral membranes in magenta (EpCAM). Scale bar = 20µm. [b]Magnified views of the confocal images. (Left) White arrows indicate the orientation of the cell-edges with respect to the radial direction. (Middle) White arrowheads highlight buckling and bending of cell edges. (Right) Red arrows indicate the orientations of the main axis of the nuclei. Scale bar= 20 µm.[c] Schematic and quantification of the tilt angle and scaled bending energy for each edge of the cortinoids shown in [b]. Arrows indicate the average direction. The center of the lumen is shown with ‘+’. The dotted curve denotes the lumen boundary. Red and blue colour scales show the tilt angle and scaled bending energy, respectively. [d] Bending energy *B* of cell edges at rest (n=20) and inflated states (n=18). ****p *<* 0.0001, Mann–Whitney U-test. [e] Tilt energy *T* of cell edges at rest (n=20) and inflated states (n=18). *p *<* 0.05, Mann–Whitney U-test. In each violin plot gray markers indicate absolute values and solid black line shows the mean value. Violin kernel bandwidth is 0.2 in all cases. Precise definitions of *B* and *T* are given in the main text.[f] Confocal snapshot of a living cortinoid in a cycle of inflation-relaxation, at radial strains epsilon =0; 0.5 and 0. Actin is shown in grayscale after contrast inversion. Magnified views of the zone marked by the red-dotted box shows “edge A” (top row) undergoing predominantly bending, and “edge B” (bottom row) undergoing predominantly tilting. Blue arrowhead mark bending of the edge and the orange arrows highlight tilting. Outlines of the bent cell edges are drawn with shade corresponding to the magnitude of the bending energy. Scale bar=20 µm. [g] Variation of the bending energy *B* and the tilt energy *T* with radial strain during inflation and deflation of edges A and B, respectively. Arrows on each curve denote the direction of change in radial strain.

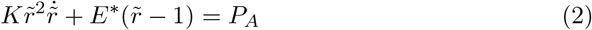

with 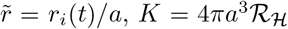 encoding an effective viscosity and 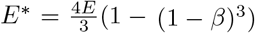 being the apparent Young modulus of the epithelium, where *β* = (*b* − *a*)*/b* is the ratio of thickness and outer radius at t=0 (*b* and *a* being the outer and inner radii at t=0, respectively). By further varying *P*_*A*_ from 1 to 4 kPa (Supp. video 6-8) and only examining the low deformation regime, fits of experimental data with the solution of Eq.2 (fig.2g) yield an Young’s modulus, *E* = 4.6 ± 1.4 kPa independent of the stage of growth after encapsulation. Additionally, this value does not vary significantly for cortinoids of different initial sizes (fig.2h). A similar elastic modulus has been numerically derived from a morpho-elastic growth model in [22]. For comparison purposes, epithelial monolayers grown on collagen layers have been reported to be as stiff as 20-30 kPa[37]. More physiologically relevant, in [38], a Young’s modulus of 20 kPa was measured for a freely suspended monolayer of MDCK cells, which is more than an order of magnitude higher than the one found here for hiPSC epithelium. On deriving the lumen pressure from equation (1), for a loading cycle, we observe minimal hysteresis between the loading-unloading pressure-strain traces (fig.2 i). The collection of all experimental data for multiple cortinoids and multiple cycles at different applied pressures shows that the rate of increase of lumen pressure decreases with increasing strain at large radial strains *>* 0.4 (fig.2 j), indicating that the epithelium is stretched without building significant stress. This shell softening signature at large strains with reversibility hints at a stress-release mechanism, which is expected to be beneficial for epiblast model growth. As a qualitative check, we perform an additional experiment in which cortinoids are allowed to grow after inflation up to 100 % strain. No significant change is observed in the growth and morphology of the cortinoids over the next 20 hours (Extended Fig. 2), showing minimal impact of the high strains.

### 2.4 Bend-tilt mode of deformation

To investigate the microscopic origin of the super-deformable behaviour, we first perform high-resolution imaging on fixed samples. To fix them in an inflated state and ensure that they have not relaxed, the cortinoids are inflated from the interior with 40% paraformaldehyde (PFA) (technical details in the Methods section) (Fig.3 a). Confocal micrographs (fig.3 a) reveal that, at the rest state, the cell edges are predominantly straight and aligned along the radial direction of the shell. The major axis of nuclei is also aligned radially, oriented towards the center (fig.3 a-b top row). At high inflation (∼ 80%), we observe a thinning of the epithelium, expected for an incompressible cell material. The nuclei are primarily aligned in the ortho-radial direction and flattened. The cell edges are either bent and/or tilted with respect to the radial direction (fig.3 a-b bottom row). To quantitatively assess the deformation of cell edges, we define (i) the tilt angle, measured between the vectors from the centre of the cortinoid to the apical and basal ends of an cell edge and (ii) the dimensionless bending energy *B*, calculated as the integral of the curvature of an edge multiplied by the arc length of the edge (see methods). A visual representation of tilt angle and bending energy is shown in (fig.3 c) on a representative cortinoid. To represent the tilt in energy terms, we also define the dimensionless tilt energy, *T* computed as the square of the previously measured tilt angle (fig.3 c and see methods). On calculating the tilt energy (fig.3 c) and bending energy (details in methods), we observe that compared to the normal state, the energies are almost twice in the inflated state (figs.3 d and e). This confirms that the edges are more bent and tilted in the inflated state. To ensure that these are not artefacts from fixation and to allow investigation of the reversibility at the microscopic level, we finally performed live confocal imaging of cortinoids during an inflation-deflation cycle (Supp. video 9) by labelling the cell edges with the SPY actin live probe (details in materials and methods). By monitoring two specific cell edges upon inflation to radial strain *ϵ* = 0.5 and deflation back to *ϵ* = 0, we observe reversible bending and tilting events (fig.3 f). The deformation traces an energetically reversible path in terms of *B* and *T* in a cycle, highlighting the elastic nature of the cell edge in both modes, i.e. bending and tilting. Taken together, the experiments described above suggest that structural deformations of the cell edges may buffer the stress in the epithelium that accounts for the softening behaviour in the stress-strain curve (Fig.2 j).

Although epithelial mechanics couples active stresses, viscous dissipation and elastic storage, the reversible response in fig.2i and fig.3g signals that we probe a elasticity-dominated limit[39]. The non-elastic terms originate from mechanochemical remodelling of the actomyosin cortex, and are governed by the turnover times of cortical actin and crosslinkers (*τ*_*o*_ ≈ 30-60 s) [40, 41] Using *ϵ*_max_ = 0.6 and *τ*_*o*_ ≈ 60 s gives a reference strain-rate of 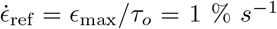. Because the strain rates in our experiments (0.6 - 1 % *s*^−1^) are at this reference rate, mechanochemical remodelling is too slow to relax or generate appreciable stress within a cycle, and cellular cortex acts like a solid yet hierarchically-flexible architecture. This elastic mechanism provides a route towards extreme deformability, complementing the active superelasticity reported earlier [42] in tissue mechanics. Whereas active superelasticity arises at low strain rates 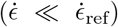, relying on actomyosin cortex remodelling, elastic superdeformability emerges at high strain rates 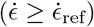.

### 2.5 Modelling the mechanical response of cortinoids

To examine how reversible elastic deformations of cell faces translate to mechanical buffering at the tissue scale, we develop a minimal model for the pressure-strain response (Supplementary sec. C). Volume-conserving inflation of an elastic cortinoid leads to in-plane biaxial extension with transverse thinning. For a continuum solid shell this necessarily entails radial compression [43]. Due to the hierarchical composite structure of epithelial tissue, an assembly of fixed-volume cells with a fluid cytosol enclosed by a thin actomyosin cortex, inflation of cortinoids can be accommodated not only by lateral face compression, as in a continuum solid, but also by the bending and tilting modes observed experimentally. Because the actomyosin cortex is thin relative to the cell size (*t*_0_ ≪ *d*), bending and tilting offer far less resistance to shell thinning than compression (*k*_bend_*/k*_comp_ ∼ (*t*_0_*/d*)^2^), and thus act as soft modes of deformation. These considerations define two limiting regimes: without soft modes, apico–basal stretching must be coupled to lateral compression; with soft modes, the effective resistance to radial compression vanishes. Fig 4b illustrates these limits for inflation of a 3D volume-conserving shell of uniform cells with faces modelled as thin incompressible Neo-Hookean sheets (see Supplementary sec. C). Removal of lateral compression causes a marked drop ∼50% in the pressure-strain relation.

**Fig. 4.**
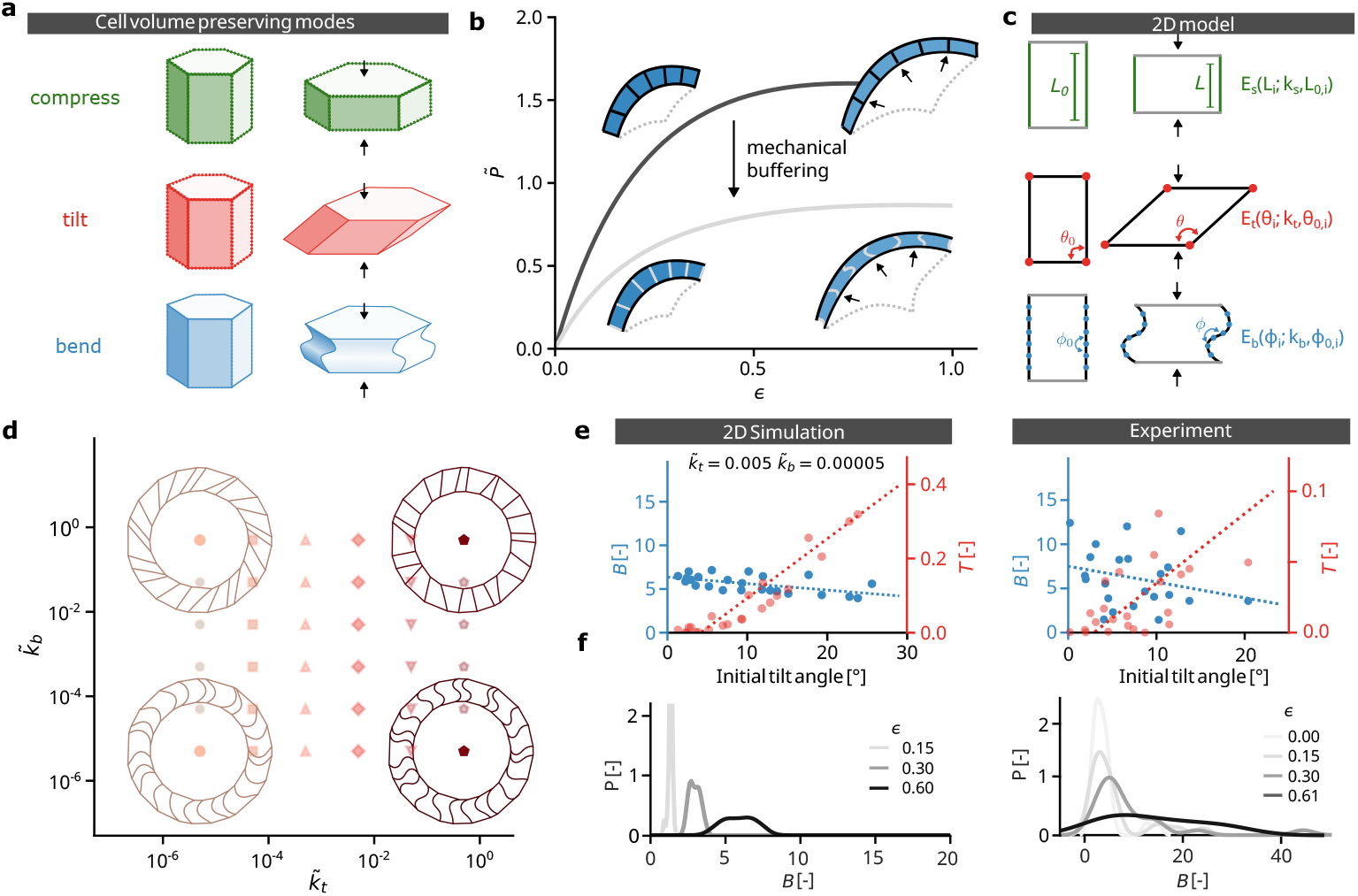
Mechanical buffering through soft modes. [a] Cell area preserving deformation modes. The compression mode requires lateral edge length change. In the tilt mode the apical and basal face move tangentially, to allow thinning of the cell without loss of lateral edge length. Similarly, the bend mode follows from buckling of the lateral face. [b] Limits for the mechanical response. Black line: apical/basal stretching and lateral compression (black line). Grey line: apical and basal stretching. The analytical pressure-strain curves describe inflation of a 3D cortinoid, assuming volume conservation of the epithelium and treating the cell faces as neo-Hookean incompressible thin sheets [see SI]. Pressure is normalized as 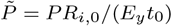. [c] 2D model for area preserving soft modes. The system incurs an energy penalty for deviations from the reference configuration, which are proportional to the stretch,tilt, and bend constants: *k*_*s*_, *k*_*t*_, and *k*_*b*_. [d] Effect of 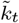 and 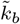 on the simulated inflation response. We display the minimum energy configuration for the limits of the explored parameter space at *ϵ* = 0.6 (solid symbols). [e] Magnitude of tilt (in light red) and bend (in dark blue) of edges at *ϵ* = 0.6 as a function of the original tilt angle for the 2D model (left) and experiment (right). Filled dots indicate experimental and simulation results. The dotted lines are linear fits to the respective data sets. For *B*, there is a weak dependence on the initial tilt angle. [f] Evolution of the bend distribution *B* for the 2D model (left) and experiment (right) during a live experiment for 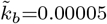 and 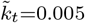.

To explore how these cell-level soft modes shape tissue mechanics between the two limits, we construct a 2D cortinoid model of a ring of cells with harmonic penalties for area change, edge stretching, whole-cell tilt, and lateral-edge bending [44–46](fig. 4a and Supplementary sec. C)). After rescaling, the parameter space reduces to two effective stiffness ratios for tilt and bend, 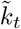 (tilt) and 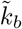 (bend). Minimizing the total energy with respect to radial strain yields the geometric evolution and the pressure-strain curve. The dominant deformation mode depends on the relative magnitudes of 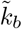 and 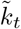: bending for 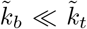, tilting for 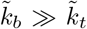, compression when 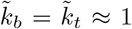, and mixed modes when 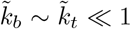 (see Supp vid 11). The resistance to soft modes tunes the mechanics: when 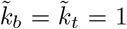; lateral compression dominates when 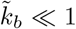 or 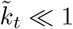, the mechanical response approaches pure apico-basal stretching, consistent with the limits of the 3D model; and for intermediate values, the mechanics interpolates between these limits (see Extended Fig. 3 and Supplementary sec. C).

The 2D minimal model links the control parameters, 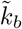 and 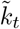, to two observables accessible in experiments: the onset strain for bending and the onset strain for tilting (extended Fig. 4 and SI). No clear onset in ⟨*B*⟩ or ⟨*T*⟩ is observed (Fig. 3g and Supp vid. 9), consistent with a much lower resistance to bending and tilting, compared to lateral compression, i.e. 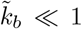 and 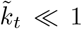. In this regime, the simulations qualitatively reproduce the experimentally observed dependence of *B* and *T* on the initial tilt angle (fig. 4d) and the broadening of the distribution in *B* (fig. 4e).

Overall, these results indicate that during rapid inflation, lateral-edge soft modes buffer the mechanics by decoupling radial thinning from the pressure-strain response.

## 3 Discussion

Our findings show that iPSC shells achieve extreme deformability by exploiting reversible soft modes of deformation. During inflation–deflation cycles, lateral cell edges of the shells which are reinforced by actin filaments bend and tilt, providing elastic pathways that buffer mechanical load by decoupling apico–basal stretching from lateral compression. These instabilities act as mechanical safety valves: they are reversible, preserve hydraulic barrier integrity, and prevent stress accumulation even at high strains.

This mode of mechanical adaptation stands in contrast with plastic or superplastic deformation in engineered materials, where large strains are achieved through irreversible processes such as dislocation motion, grain boundary sliding or other energy-dissipating [47–49]. iPSC shells instead occupy a distinct regime, where elastic buffering through reversible instabilities enables large shape changes without permanent damage. In this sense, they demonstrate that super-deformability can emerge not only from softness or plasticity but also from the activation of alternative, low-energy structural modes.

The superdeformability of iPSC shells is thus best understood as the outcome of multiscale mechanics. The extreme softness of the epithelium is a consequence of its meso-scopic structure, wherein the cytoskeleton provides access to reversible bending and tilting instabilities that accommodate thinning and radial expansion leading to large, reversible deformations while preserving epithelial architecture.

Such a strategy may be particularly valuable in morphogenesis, where epithelia must often undergo extreme shape changes while maintaining functional integrity. The ability to buffer stress through tilt and bend instabilities provides robustness against rupture, offering a physical mechanism for the complex morphologies seen during early embryogenesis. More broadly, our results suggest that developmental tissues may harness reversible instabilities as elastic reservoirs, complementing or replacing plastic remodeling, to ensure both mechanical resilience and morphogenetic flexibility.

## 4 Methods

### 4.1 iPSC cell culture and encapsulation

hiPS cells purchased from Corriell Institute (AICS-0023) are cultured using standard protocols in T25 plastic flasks (Corning-Ref: 353109), coated with Matrigel (Corning-Ref: 354234). Once at a confluency of 75-80 %, the cells are detached using Acutase (Stemcells-Ref: 07920) and resuspended in culture media at the required concentrations. The culture medium used for cell and capsules culture is mTeSR™ Plus (Stemcells-Ref: 100-0276).

As previously described [11, 16], we used the Cellular Capsule Technology (CCT) to produce capsules of an alginate shell (AGI, I3G80), enclosing 7-10 hIPSCs in an extra-cellular matrix, Matrigel (Corning-Ref: 354234). Equal volumes of a diluted cell solution of 0.8-1M cells/ml and Matrigel solution are used as a cell solution mixture. Briefly, using a microfluidic chip, a co-axial compound jet of an outer envelope of alginate 2 % [w/v] and the cell solution, separated by an intermediate solution of D-sorbitol (300mM, 85529, Sigma), is produced. The jet is allowed to fall freely and break into core-shell droplets that are collected in a calcium chloride bath (100mM, 7774-34-7, Themo scientific), which leads to gelification of the alginate and formation of the capsules. The capsules of hiPSCs are rinsed immediately with DMEM (Biowest L0102) to remove excess calcium and cultured in mTeSR™ Plus culture medium at 37°C and 5 % *CO*_2_. ROCK inhibitor Y-27632 dihydrochloride (Tocris-Ref: 1254/10) at 10 mM is added and replenished at 1/1000 dilution every day for the first 48 hours after encapsulation. After 48 hours (day 2), culture media with Penicillin-streptomycin (Gibco™ 15140122) in 1:100 dilution is used and replaced every 48 hours.

### 4.2 The nano-inflation apparatus

The nano-inflator comprises of a pneumatic pressure source connected to a pipette mounted on a micro-manipulator.

Pipette fabrication: Capillaries (TW100-6 WPI, *d*_*i*_=0.75 mm, *d*_*i*_=1 mm)) are shaped using a pipette puller (P-1000, Sutter Instruments) and clipped with a microforge (MF2, Narishige) to the required diameters for the experiments (2-3 µm). The pipette needs to be stiff enough to penetrate the capsule and the cell layer without bending. Increasing the pipette diameter enhances the stiffness but can cause substantial damage to the cell layer, potentially leading to leakage during inflation. Hence, an empirically optimized pipette profile is designed using a customized program on the pipette puller. The pipette is pulled through a two-step process of heating and pulling (parameters: Ramp: 525, 1-Heat: 525, Pull: 0, Vel: 20, Time: 200, Pressure: 500, 2-Heat: 535, Pull: 90, Vel: 75, Time: 200, Pressure: 500,. Using a premounted scale on the microforge objective, the pipette is positioned precisely and hot-cut at the desired diameter.

Measurement of pipette diameter: The hydraulic resistance ℛ_*H*_ of the microfluidic circuit is dominated by the resistance of the pipette tip since the hydraulic resistance, 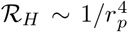. As a consequence, the tip must be measured accurately. Two different techniques are used. To mitigate the aberrations caused by the cylindrical shape of the glass micrometric pipette tip, the pipette is dipped in water and imaged with a high magnification objective (Nikon CFI Plan Apochromat Lambda D 40X, NA-0.95, WD-0.21) in an inverted microscope (Nikon Eclipse Ti2). The other technique involves the global shape of the pipette. The end of the pipette is conical: the inner diameter increases linearly from the tip of the pipette over a length of at least 1 mm. Instead of measuring the diameter at the fine tip, the diameter is measured with high precision at 0.5-1 mm away from the tip using a 20X, 0.8 NA (CFI Plan Apochromat Lambda D 20X, NA-0.8, WD-0.8) objective and linearly extrapolated to obtain the diameter at the tip. A water soluble dye is used to fill up the pipette to further enhance the contrast and accuracy of the measurements.

Pipette filling: In most inflation experiments, the pipette is filled with an osmotically balanced liquid composed of D-sorbitol (300mM, 85529, Sigma) and traces of a dye, Napthol Blue-Black (2% [w/v] Sigma 195243), to enhance optical contrast for further image analysis and shell rupture detection. The solution was preliminarily sterile filtered with a 0.22 µm filter to remove impurities and debris that could clog the pipette. The pipettes are back filled with a microneedle (MicroFil flexible needle, WPI MF28G-5) and mounted on the sleeve of the head of the pipette holder in the micromanipulator (Eppendorf InjectMan 4). In the holder head, a rubber sleeve tightly grips the pipette and care is taken to fill the small void behind this sleeve with an excess of the injected fluid to ensure no bubbles.

Monitoring flow rate: The fluid circuit from the reservoir to the pipette holder consists of a tubing with an inner diameter of 1 mm. A flow-meter (Sensirion LG16-0430D) was first connected to the line to measure the flow rate in time. The reservoir is positioned slightly below the vertical level of the pipette outlet to create a backflow (from the pipette to the reservoir) due to hydrostatic pressure. This ensures that during experiments, positive pressure is needed to reach a state of no-flow in the system and hence avoids the risk of uncontrolled flow in the system due to hydrostatic pressure. To pressurize the system, a pneumatic pressure source (Biophysical tools) with a resolution of ∼0.05 mbars with a range of 500 mbars (5 kPa) is used. When a shell of initial lumen volume 0.2 nL needs to be inflated by 10% in radius, or 30% in volume, given that the accuracy of the flow-meters is 10^−1^ nL/s, the experiment would be over in a fraction of second, making the analysis inaccurate due to a lack of temporal resolution. Instead, we directly monitored the increase of lumen volume. The flowmeter is only used to check for bubbles in the fluidic line.

### 4.3 Immunostaining

Capsules were collected and fixed with 4% Paraformaldehyde (PFA) (Fisher F/1501/PB17) on days 2-6 following encapsulation. Counterstaining was performed with Hoechst 33342 (Life Technologies H3570) and Phalloidin (A12379, A12380, Invitrogen) (fig.1b). The hiPSCs TJP1 are endogenously tagged with mEGFP. To image cortinoids in the inflated state, 40 % paraformaldehyde is injected to fix them from the inside. Once a shell is inflated, it is immediately pipetted from the dish and transferred into a bath of 4 % PFA and 0.2% glutaraldehyde (Sigma G6403). The samples were kept in this solution for 2 hours, before washing with DMEM. To image shells with highly resolved labelling of the cell edges, immunostaining was performed. Permeabilization and blocking prior to immunostaining were performed at room temperature for 2 hours in a solution of DMEM (without red phenol, 11054020, Gibco), Triton X100 0.1% (A4975, AppliChem Panreac), Tween 0.2% (Sigma P1397) and FBS 10% (Capricorn Scientfic FBS-16A). To stain the baso-lateral edges of the cells in the epithelium (fig.3a), primary antibody epCAM (EpCAM Monoclonal MA5-12436, Thermofisher) was used at 1:100 dilution, followed by a conjugation with a secondary antibody (Alexa-fluor 647 A32787, Thermofisher) at 1:500 dilution. Primary antibodies were incubated for 72h, and secondary antibodies were incubated for 24h following thorough rinsing in DMEM with 1% FBS.

### 4.4 Imaging

Live monitoring of hiPSC growth was performed with an inverted microscope (NIKON, AZ100) in a bright-field mode of acquisition with a macro-objective (Nikon AZ plan fluor 5X). Immunofluorescent images (fig.1b) were imaged in 3D with a spinning-disk microscope Leica DMI8 (Leica Microsystems) equipped with a confocal Scanner Unit CSU-W1 T2 (Yokogawa Electric Corporation) using a 20X multi-immersion objective at Bordeaux Imaging Center. For high optical sectioning of the inflated fixed samples(fig.3a), a scanning-confocal microscope (Leica DMI6000 TCS SP8 X) with a 20X/0.75 immersion objective was used to image the samples in 3D.

Inflation experiments with black dye in bright field mode has been performed on an inverted microscope (Nikon Eclipse Ti2) with a 20X objective (Nikon CFI S Plan Fluor LWD ADM 20XC NA-0.7 WD-2.3) Inflation of hiPSCs cortinoids with fluorescent dye (fig.2 a) was carried out on the same microscope with a spinning head (Cicero CrestOptics, Imagxcell) with the same objective. To stain hiPSCs shells in live, non-toxic SPY-Actin 555 (actin) and SPY-DNA 650 (nuclei) (Spirochrome) are added to the culture media at 1:500 dilution, and incubated at 37°C, 5% *CO*_2_ for 1 hour inside the incubator. For live imaging of the inflations (fig.3 f), selected cortinoids are seeded on a glass bottom Petri dish and placed inside a microscope stage incubator (Okolabs H301-K). Imaging is performed with a scanning confocal microscope (Leica Stellaris DMI 8) using a 25X water immersion obejctive (Leica HC FLUOTAR L VISIR 25x/0.95 WATER). To inflate the shells in this configuration, a small customized manual stage fitted with XYZ micrometers and a pipette holder is used. The same fluidic circuit and glass micropipette as discussed above are used.

### 4.5 Extracting morphological parameters

Similar to the image analysis methods described in [21], a image analysis pipeline using ImageJ and MATLAB2023 (MathWorks) was developed to track the inner radius and outer radius of the cell layer. Adding a dye during the inflation experiment facilitates the visualization and delineation of the inner radius of the cell layer. Briefly, using gradients of intensity in local windows of 20 pixels across the cell layer, the inner and outer radius are determined at different angular positions of the shell. The region in the vicinity of the pipette is discarded for quantification. The inner and outer position of the cell layer are measured every 5° around the shell, and the mean value is used to compute the lumen radius and thickness of the shell at each time point (Supp Video 10). The standard deviation of these measurements takes into account the non-circularity of the shell. In the case of the data processed for Fig.1 (b), analysis of the shell volume is carried out manually by drawing masks over the lumen and outer edge of the shell to extract the radii derived from the mean of the major and minor axes of an elliptical fit over the luminal and external masks.

To derive the lumen pressure (fig.1 g), an exponential decay is fitted to the first 10 minutes of the variation of lumen volume with time. The flow rate at the instant of inserting the pipette is extracted from the initial slope of this fit. This flow rate is multiplied by the hydraulic resistance of the fluid circuit (see above) to measure the lumen pressure in the rest state.

To trace the shape of the cell edges as the shell is inflated (fig.3 d), free hand lines are drawn on the edges of the cells at different strains. Twenty equally spaced interpolated points are used to discretize each edge at each time point. This removes problems of pixel-fractality that may arise on free hand drawing. The bending energy is calculated from the integration of the square of the curvature (*κ*) over the arc length (L) of each edge, *B* = *L* ∫ *κ*^2^*ds. B* here is non-dimensional and is proportional to the bending energy density (see Supplementary sec D.3). The tilt energy is taken as the square of the angle between the apico-central and baso-central vectors, *T* = *θ*^2^. Average values of these scaled bending and tilting energy are represented as ⟨*B*⟩ and ⟨*T* ⟩, respectively.

## Supplementary information

All supplementary data has been included in a separate document.

## Supplementary videos

1. Supplementary Video 1: Growth of hiPSC shells cultured in Matrigel laden alginate capsules from Day 4 (t=0 h) to Day 6.
2. Supplementary Video 2: Micropipette inserted in the lumen of a cortinoid to pressure the lumen pressure. The pipette is withdrawn after 15 minutes and the shell is further monitored. Refer to Extended fig. 1.
3. Supplementary Video 3: Collection of shells with the pipette inserted and removed for lumen pressure quantification.
4. Supplementary Video 4: Inflation of a shell with fluorescent dextran (FITC-Dextran 20 kDa at 0.1 %) shown in bright field (left) and in fluorescence (right). ‘ON’ on the video denotes the time when a constant pressure step is applied.
5. Supplementary Video 5: Multiple inflation and relaxation cycles on a shell at a constant pressure step of 1kPa. ‘ON’ on the video denotes the time when a pressure step is applied.
6. Supplementary Video 6: shell inflation at a constant pressure step of 1kPA.
7. Supplementary Video 7: shell inflation at a constant pressure step of 2kPA.
8. Supplementary Video 8: Cortinoid inflation at a constant pressure step of 4kPA.
9. Supplementary Video 9: Live inflation and relaxation of a shell at a constant pressure. (left) Bright field acquisition. (right) Actin of the cells marked with fluorescent SPY-actin is shown in grayscale.
10. Supplementary Video 10: Automated quantification of lumen radius, thickness and outer radius of a shell undergoing an inflation process at a cosntant pressure step of 1kPa. The detection of the thickness of the cell layer at each angular position is shown in green. The center is marked with a ‘+’.
11. Supplementary Video 11: Limits of elastic deformation of 2D cortinoids for 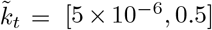 and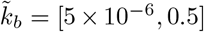, Including the compression mode (top right) and the two soft modes: tilt (top left) and bend (bottom right).

## Acknowledgements

We thank Jacques Leng and Jean-Baptiste Salmon for fruitful initial discussions. The authors also thank the BiOf (BioImaging and Optofluidics) team for everyday support. The authors thank the engineers for their assistance in the imaging experiments the Bordeaux Imaging Centre, a service unit of the CNRS-INSERM and Bordeaux University (ANR-10-INBS-04).

## Declarations

### Funding

AJ acknowledges the Agence Nationale Recherche Technologie (ANRT) for PhD fellowship. PN was supported by the French National Agency for Research (ANR-21-CE18-0038; ANR-23-CE45-0016) and from the Institut National du Cancer (PLBIO23-097). GR was supported by the French National Agency for Research (ANR-21-CE45-0028). This study also received financial support from the French government in the framework of the University of Bordeaux’s France 2030 program/GPR LIGHT. J.T. was supported by an EMBO Postdoctoral Fellowship (ALTF 2022-7). LM thanks the Simons Foundation and the Henri Seydoux Fund for partial financial support.

### Conflict of interest/Competing interests

The authors declare no competing interests

### Ethics approval and consent to participate

NA

### Consent for publication

Yes

### Data availability

All data are available here: zenodo/Reversible superdeformability of hiPSCs cortinoids

### Materials availability

NA

### Code availability

Code related to experiments is available at- github.com/BiOflab/hiPSC-superdeformability. Code related to simulations and theory is available at-github.com/mahadevan-harvard/hiPSC-superdeformability

### Author contribution

PN, KA, MF and AJ conceived and designed the project. AJ performed all the experiments and analyzed the results. AJ with BG performed live imaging. GR advised AJ and gave regular feedback on the results. JT and LM developed theoretical frameworks. JT performed all the simulations and model calculations. AJ, JT, PN, and LM discussed the results, contributed to the mechanistic understanding and co-wrote the manuscript.

## Extended figures

**Extended Figure 1.**
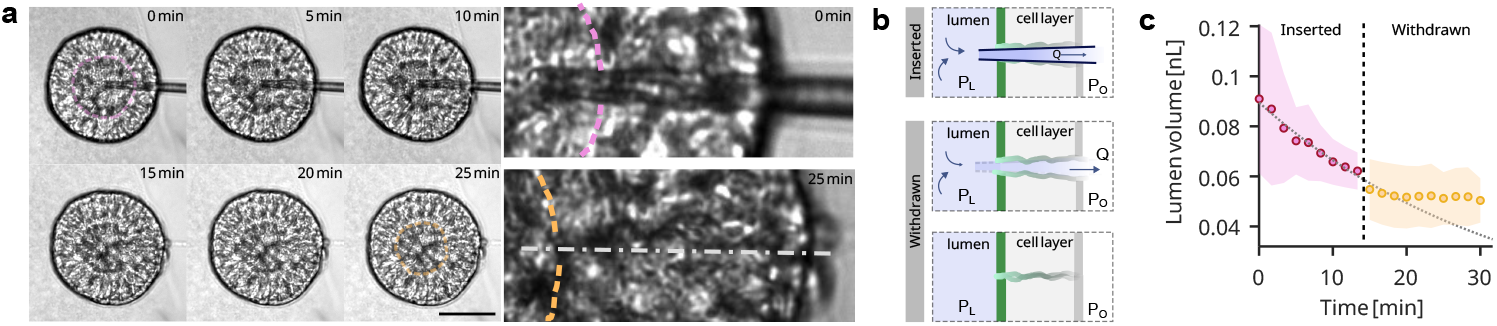
Lumen drainage before and after insertion of pipette. [a] (Left) Time-lapse of a lumen with a pipette inserted and withdrawn. The lumen at t=0 min is outlined with a pink dotted curve and at t=25 min is outlined with orange dotted curve. (Right) Zoomed in view of the region of the shell in which the pipette is inserted (top) at t=0 min and (bottom) after removal at t=25 min. The outlines of the lumens are shown in pink (top) and orange (bottom) dotted curves. The axial line in the bottom shows the midline of the pipette that was removed. [b] Schematic showing the insertion, drainage and healing of the cell layer on removing the pipette. [c] Variation of lumen volume with time on inserting the pipette and withdrawing the pipette (shown in pink and yellow. The dotted curve is an exponential fit to the points in the inserted region.

**Extended Figure 2.**
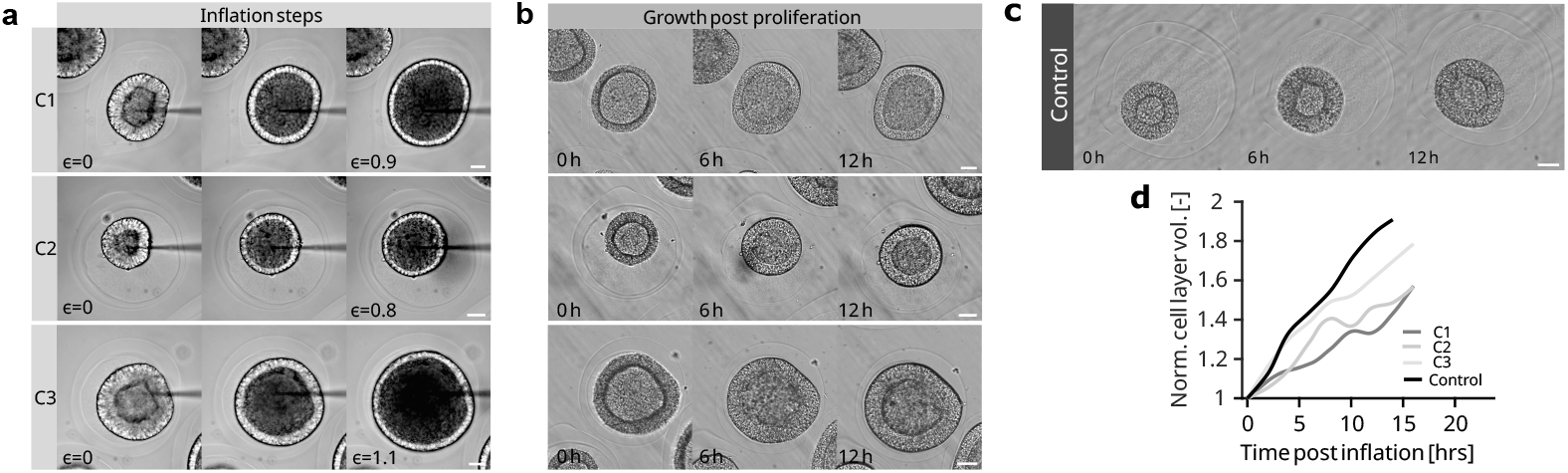
shell dynamics before and after inflation. [a] Inflation of shells at constant pressure upto shown strains. In case of C2, the shell ruptures. [b] snapshots of of shell proliferation post inflation. t=0h is 1 hour post the inflation. [c] Snapshots of a shell growing normally without any inflation. [d] Variation of cell layer volume rescaled with the initial volume of the cell layer at t=0 with time post inflation.

**Extended Figure 3.**
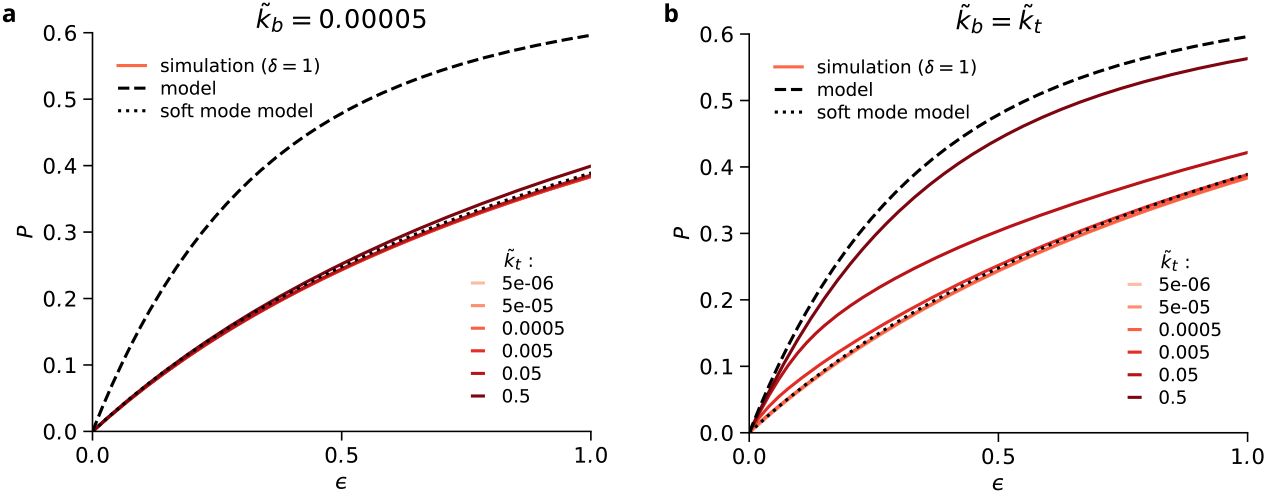
Mechanical response of the 2D shell model. Pressure-strain curves for varying tilt resistances 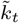 (legend). For all simulations we consider disorder *δ* = 1. [a] 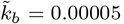, and [b] 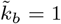1. In both panels, the stretched based limit (black dashed) and soft mode limit (black dotted) from Eq.D43 (supplementary sec.D)are shown for comparison.

**Extended Figure 4.**
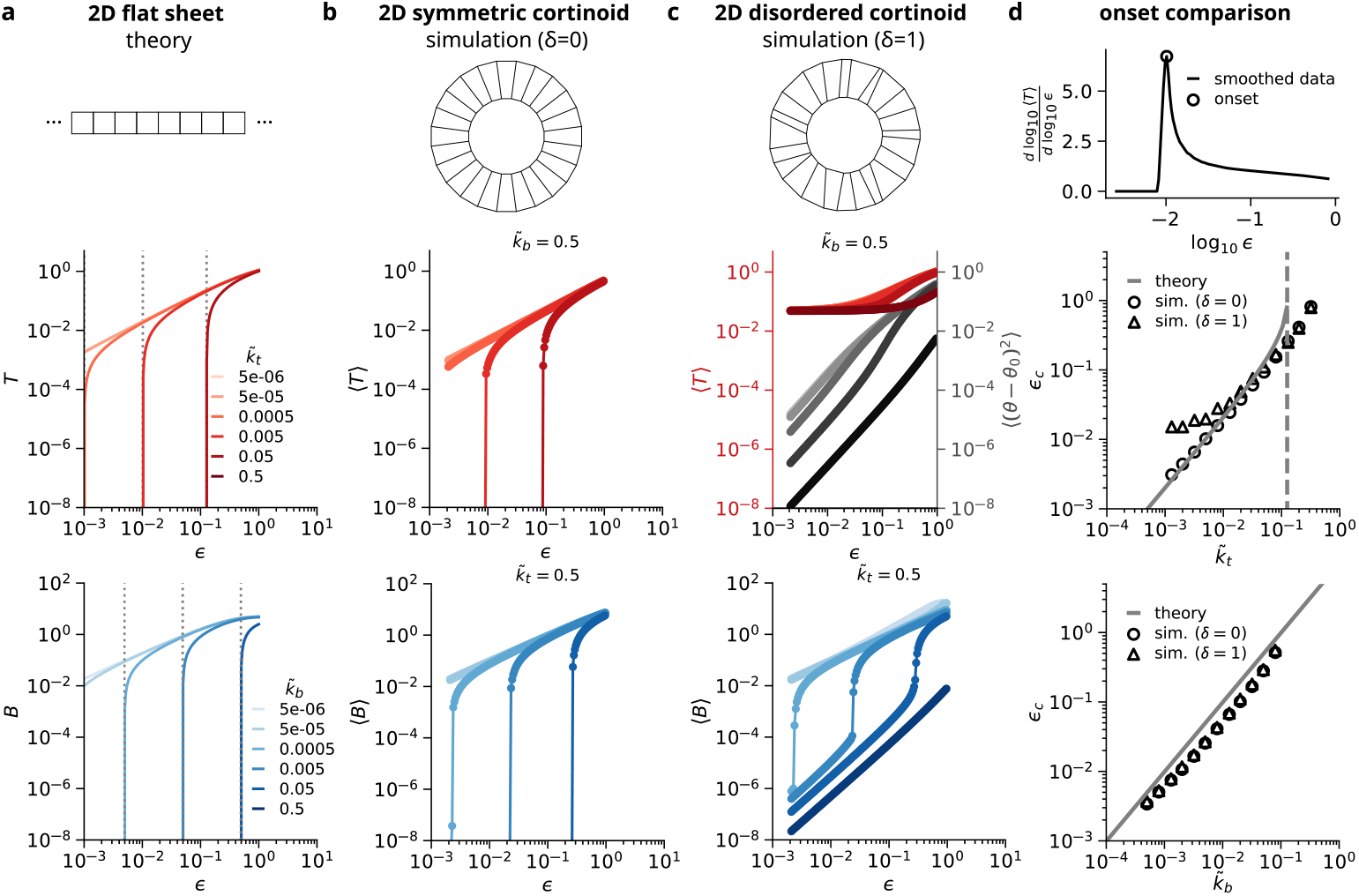
Development of soft modes versus strain. [a] 2D flat sheet model, [b] 2D symmetric cortinoid model (*δ* = 0), and [c] 2D disordered cortinoid model (*δ* = 1). [d] Onset strain *ϵ*_*c*_ from [a-c]. Top row (tilt *T*): colors correspond to 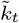. In [a] we plot the numerical solutions of Eq.D48with predicted onset from Eq.D49(grey dotted). Panels [b,c] show simulations (Sec. D1) with 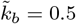 to suppress the bending mode. In [c] we also plot ⟨(*θ* − *θ*_0_)^2^⟩ (grey; intensity increases with 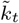) Bottom row (bend *B*: colors correspond to 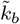. In [a] we plot numerical solutions of Eq.D59as well as the predicted onset from Eq.D60on the linearized solution (black dotted). Panels [b,c] show simulation results (Sec. D1) with 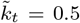 to suppress the tilt mode. In [d], *ϵ*_*c*_ is extracted as the strain at the maximum slope of *log*_10_ ⟨*T* ⟩ or *log*_10_ ⟨*B*⟩ versus *log*_10_ *ϵ*. Prior to differentiation the data are smoothed using a 5 point moving average. An example is shown for 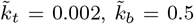. The onset for tilt with *δ* = 1 is determined based on ⟨(*θ* − *θ*_0_)^2^⟩.

## Notes

https://zenodo.org/records/17153949?preview=1&token=eyJhbGciOiJIUzUxMiJ9.eyJpZCI6ImI0YjdjZmY4LWI0MGUtNGRlYy1iZWNmLTc4M2VmOTkxNmVlNyIsImRhdGEiOnt9LCJyYW5kb20iOiI4Yjg2ZWMxYmY5M2E3MTY0OWRlYTIxNTJiYWE1NTM2ZSJ9.dSWgD7UVzyigH9BNJBWfty0lDmveI31JuS2ZTRYtu0VPZAn04WIayFI9jXECQugJyxPG9kiwNM2J2VISQppbFA

### Competing Interest Statement

The authors have declared no competing interest.

