## Supplementary file for "Reversible superdeformability of hiPSC epithelial cortinoids"

#### A Calculation of $\mathcal{R}_{\mathcal{H}}$ and lumen pressure

To calculate the hydraulic resistance used in the estimation of lumen pressure, we measured the pipette profile in the vicinity of the tip. From the measurements, we extract the tip diameter (fig.A1 a-c) and the taper angle. The taper of the pipette $\alpha$ is determined to be 0.621° and this profile extends to at least 500 µm from the tip (fig.A1 a). Using the Hagen-Poiseuille equation for a conical cross-section, we derive the $\mathcal{R}_{\mathcal{H}}$ from equations (A1 and A2). Since $\mathcal{R}_H \sim 1/{d_p}^4$, the resistance of the pipette from the tip upto $L_c$ dominates the total resistance of the fluidic circuit (fig.A1 d). Resistance increases by less than 5 % beyond 240 µms. Hence, precise measurements of the pipette shape up to this length is sufficient to estimate the hydraulic resistance of the entire fluid circuit. To extract the elastic modulus (E) in the small strain regime, the solution of equation(2) (refer to main text) is fit with the experimental data of strain versus time. Here E is the only unknown, but the $\mathcal{R}_{\mathcal{H}}$ in equation (2) is allowed to vary within the limits of the experimentally measured variation to ensure better fits. From these optimized $\mathcal{R}_{\mathcal{H}}$ values, the tip diameter is back calculated to compare with the measured pipette diameters and we observe a good match within the limits of error in measurements of the pipette diameter. (Fig.A1 e).

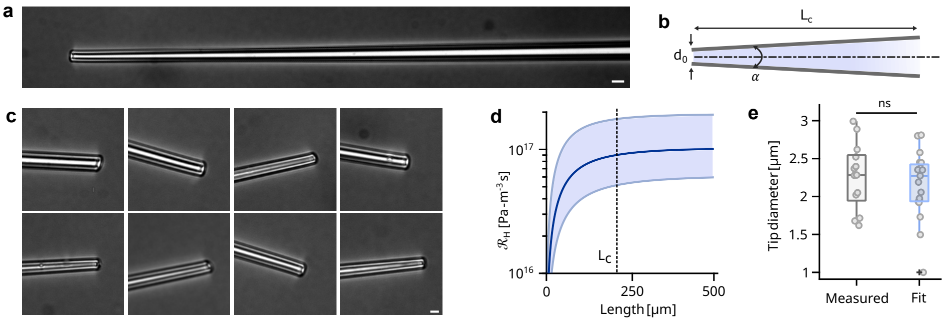


**Fig. A1** **Quantification of $\mathcal{R}_{\mathcal{H}}$.** [a] Bright field image of a micropipette from the tip to a length of 360 µm. Note the taper in the pipette. Scale bar = 10µm.[b] Schematic of the pipette highlighting the tip diameter $d_0$, taper angle $\alpha$ and the critical length $L_c$. [c] Zoomed-in snippets of the pipette tip after manufacturing. Scale bar = 5µm. [d] Variation of the hydraulic resistance $\mathcal{R}_{\mathcal{H}}$ for $d_0 = 2.3 \pm 0.43$ µm over the length of the pipette measured from the tip (data taken from the optimized fits, n= 19). $L_c$ is the critical length beyond which there is negligible increase ($< 5\%$) in $\mathcal{R}_{\mathcal{H}}$. [e] Tip diameter measured directly from images and tip diameters obtained from the optimized fits of eq. 2 in section 2.3. Box plots show the median (dark line), 25th and 75th quartile (the box), maximum and minimum (extent of vertical lines) and outliers (shown as '+'). p-value = 0.51 from standard independent t-test.

$$\mathcal{R}_{\mathcal{H}} = \int_0^L \frac{128\mu}{\pi d_p(l)^4} dl \tag{A1}$$

$$d_p(l) = d_0 + 2l \tan(\alpha/2) \tag{A2}$$

### B Continuum models for the mechanical response of cortinoids

At the coarsest level, the epithelium can be modelled as a continuum material, allowing the use of classical elasticity theory to describe cortinoid inflation. In this framework, the inflation pressure is related to a linear modulus $E$, the initial inner radius $R_{i,0}$ and the initial thickness $H_{i,0}$. A common simplification is to assume that the hoop stress is uniform across the shell: the thin shell approximation. However, this assumption breaks down for thick shells ($H_{i,0} \sim R_{i,0}$) where the hoop stress $\sigma_{\theta\theta}$ varies considerably along the radial direction. Below we first describe how we estimate the linear modulus of cortinoids by combining a model for unsteady inflation with an analytical expression for thick shells at small strains. We then show how a neo-Hookean thick shell can be formulated in terms of our system to interpret the large-strain regime. We close with an interpretation of experimental results.

#### B.1 Unsteady thick-shell elastic model at low strains

To estimate the pressure inside the lumen during the inflation process at a constant pressure of $P_A$, we use the radial displacement of an elastic thick shell subjected to an unsteady internal pressure.

$$P_L(t) = P_A - 4\pi r^2(t)\dot{r}\mathcal{R}_H \tag{B3}$$

$$u_r\left(r_i\right) = \frac{3\left(p_i - p_o\right) r_i r_o^3}{4E\left(r_o^3 - r_i^3\right)} \tag{B4}$$

where $r_i$ and $r_o$ are the inner and the outer radius of the shell. For small displacements when the inner radius, $r_i = a$, we approximate $u_r\left(r_i\right) \approx r_i(t) - a$. Since there is no external pressure, $p_0 = 0$ and pressure inside the lumen at a given time is $p_i = P_A - 4\pi r(t)^2\dot{r}\mathcal{R}_{\mathcal{H}}$. The outer radius is $b = a + h$, where $h$ is the thickness of the shell. Normalizing $r_i(t)$ with the initial lumen radius, $a$ and using $\tilde{r_i} = r_i(t)/a$, the lumen radius can be explicitly expressed as a function of time in the following way.

$$\tilde{r_i} - 1 = \frac{3b^3}{4E(b^3 - a^3)}\left(P_A - 4\pi a^3\tilde{r}^2\dot{\tilde{r}}\mathcal{R}_{\mathcal{H}}\right) \tag{B5}$$

$$K\tilde{r}^2\dot{\tilde{r}} + E^*(\tilde{r} - 1) = P_A \tag{B6}$$

where $K = 4\pi a^3\mathcal{R}_{\mathcal{H}}$ and $E^* = \frac{4E}{3}(1 - (1-\beta)^3)$ with $\beta = (b-a)/b$.

#### B.2 Thick-shell elastic model at high strains

Although no closed-form expression exists for the pressure-strain relation in thick shells up to large strains, it can be computed numerically as detailed below.

Exploiting the spherical symmetry, we simplify the problem by working in spherical coordinates ($r$, $\theta$, $\phi$). Under symmetry, the stress tensor satisfies $\sigma_{\theta\theta} = \sigma_{\phi\phi}$ and $\sigma_{r\theta} = \sigma_{r\phi} = \sigma_{\theta\phi} = 0$, reducing the problem to a one-dimensional radial system.

For incompressible materials, the strain field is fully determined by geometry. As a result, we can describe how a material point at a radial position $R$ in the reference configuration maps to its deformed radial coordinate $r$, as a function of the undeformed inner radius $R_i$ and the lumen stretch $\lambda_i = r_i/R_i$ (or strain $\varepsilon_i = \lambda_i - 1$):

$$r(R) = (R^3 + R_i^3(\lambda_i^3 - 1))^{1/3} \tag{B7}$$

The hoop stretch is then given by $\lambda_{\theta\theta} = \lambda_{\phi\phi} = r/R$. From the incompressibility condition $\lambda_{rr}\lambda_{\theta\theta}^2 = 1$ it follows that the radial stretch is $\lambda_{rr} = (R/r)^2$.

To compute the internal pressure, we combine the strain profile with the material properties to obtain the Cauchy stress tensor $\boldsymbol{\sigma}$. For incompressible materials, the total Cauchy stress is given by

$$\boldsymbol{\sigma} = \boldsymbol{\sigma}' - p\boldsymbol{I} \tag{B8}$$

where $p$ is not simply the mean (hydrostatic) stress, but a Lagrange multiplier field introduced to enforce the incompressibility constraint $\lambda_{rr}\lambda_{\theta\theta}^2 = 1$. Because of this, we cannot compute the internal pressure directly from the boundary stretches. Instead we must consider how the stress varies throughout the shell, as shown below.

To find the pressure differential across the shell, we begin with the fact that all internal forces must be balanced. Since pressure acts along the radial direction, we consider the radial component of the force balance $\nabla \cdot \boldsymbol{\sigma} = 0$ in spherical coordinates:

$$\frac{d\sigma_{rr}}{dr} + \frac{2}{r}(\sigma_{rr} - \sigma_{\theta\theta}) = 0\ . \tag{B9}$$

This relation must hold throughout the shell, including at the boundaries. We rewrite and integrate both sides:

$$\int_{r_o}^{r_i} \frac{d\sigma_{rr}}{dr}\, dr = -\int_{r_o}^{r_i} \frac{2}{r}(\sigma_{rr} - \sigma_{\theta\theta})\, dr\ . \tag{B10}$$

Evaluating the left-hand side gives $\sigma_{rr}(r_i) - \sigma_{rr}(r_o)$. Assuming the outer surface is traction-free, we have $\sigma_{rr}(r_0) = 0$, so the right-hand side yields $\sigma_{rr}(r_i)$, which must balance the internal pressure, so that $\sigma_{rr}(r_i) = -P$. Therefore,

$$P = -\sigma_{rr}(r_i) = \int_{r_o}^{r_i} \frac{2}{r}(\sigma_{rr} - \sigma_{\theta\theta})dr\ . \tag{B11}$$

The exact form of the stress difference $\sigma_{rr} - \sigma_{\theta\theta}$ inside the integral depends on the constitutive model (see below), but since it is uniquely determined by the material parameters (e.g., Young's Modulus $E$) and the known stretch field, the internal pressure $P$ can be computed via numerical integration. For an incompressible Neo-Hookean material, the Cauchy stress in each principal direction is given by

$$\sigma_{rr} = \mu\lambda_{rr}^2 - p \tag{B12}$$

$$\sigma_{\theta\theta} = \mu\lambda_{\theta\theta}^2 - p \tag{B13}$$

with shear modulus $\mu = E/3$ in the incompressible limit. As before, the isotropic Lagrange multiplier field $p$ cancels when taking the stress difference:

$$\sigma_{rr} - \sigma_{\theta\theta} = \mu(\lambda_{rr}^2 - \lambda_{\theta\theta}^2) \tag{B14}$$

For infinitesimal inflation of an incompressible shell, Eq. B7 can be approximated as $r(R) \approx R + \epsilon_i R_i^3/R^2$. Substituting this into Eq. B11 and keeping terms to first order in $\epsilon_i$ yields

$$P \approx \frac{4E}{3}\left[1 - (1-\beta)^3\right]\epsilon \tag{B15}$$

so that the slope is equal to the effective modulus $E^*$ we used in the unsteady model in Appendix B1.

##### B.3 Comparison with experiment and assumptions

In the thick-shell model we approximate the epithelial sheet as a continuum material. This approach allows a coarse-level comparison with other tissue types even if their mechanics is probed in a different geometry.

For example, we can use the model to extract a linear modulus of $E = 4.6 \pm 1.4$ kPa from our experiments on hiPSC shells. Uniaxial extension tests on MDCK cells [1] suggest a significantly lower value in the linear regime (low strains). This might be explained by the less pronounced actin belts in MDCK cells.

We also note that our experiments do not provide evidence for strain-stiffening behaviour as emerges in uniaxial extension of MDCK cells [1]. This difference likely results from the geometry of our experiments. Firstly, the area growth and thinning of the tissue upon inflation leads to a softening response in the pressure strain curve, potentially overshadowing the non-linear strain-stiffening expected for actin filament networks [2]. Secondly, the high $\beta$ values (thick-shell regime) imply strong strain gradients along the radial direction, so that a majority of the strain is localized near the apical face. In contrast to uniaxial extension, where strain is approximately homogeneous throughout the thickness of the epithelial sheet. This could mean that other cell-components that are known to contribute the cell mechanics at intermediate to high strains, such as intermediate filament networks [3–5], are less important in our experiments.

#### C Cellular models for the mechanical response of cortinoids

Epithelia are hierarchical composite materials. At the cellular level a minimal description is a fluid cytosol enclosed by a thin actomyosin cortex. The mechanics of a cortinoid can therefore be approximated by considering the deformation of the apical, basal and lateral cortical faces. Below we consider a model for inflation in two limiting cases: one in which the lateral faces undergo radial compression, and one in which their deformation is dominated by soft modes. We close this section with an interpretation of experimental results.

#### C.1 Cellular thin-sheet model

***Strain energy functions***

We write the total strain energy as the sum of the contributions from the apical, the basal, and lateral faces:

$$E_{\text{tot}} = E_{\text{apical}} + E_{\text{basal}} + E_{\text{lateral}} \quad . \tag{C16}$$

We assume resistance to deformation is dominated by the actomyosin cortex, and approximate it as a thin incompressible elastic sheet. For a neo-Hookean incompressible solid the strain energy density is

$$W = \frac{\mu}{2}(\lambda_1^2 + \lambda_2^2 + \lambda_3^2 - 3), \qquad \mu = E_y/3, \tag{C17}$$

with $\lambda_i$ the principal stretches. Multiplying by the reference volume of a face, $V_0 = t_0 A_0$ ( $t_0$ cortex thickness, $A_0$ undeformed surface area), gives the strain energy function. Incompressibility dictates $\lambda_1\lambda_2\lambda_3 = 1$. For the inflation of the apical and basal faces, we can write this condition in spherical coordinates as $\lambda_\theta\lambda_\phi\lambda_t = 1$. Symmetry gives $\lambda_\theta = \lambda_\phi = \lambda$, so that $\lambda_t = 1/\lambda^2$, yielding,

$$E_{\text{apical/basal}} = \frac{1}{6}E_y t_0 A_0 \left(2\lambda^2 + \lambda^{-4} - 3\right) \quad . \tag{C18}$$

For the lateral faces, inflation leads to tangential stretch and radial compression. To write the strain energy function, we approximate each cell as an $n$-gonal frustum of volume $V_{\text{cell}} = V_{\text{shell}}/N_{\text{cells}}$, with reference height $H_0 = R_{o,0} - R_{i,0}$. At radius $R_0 \in [R_{i,0}, R_{i,o}]$, the cross-sectional area scales as $R_0^2$, so its circumference is $C_0(R_0) = \alpha_n 2\sqrt{4\pi R_0^2/N_{\text{cells}}}$, with $\alpha_n = \sqrt{n\tan(\pi/n)/\pi}$ the polygon-to-circle perimeter factor ($\alpha_n = 1$ for circular faces). From the circumferential stretch $\lambda_C(R_0)$, the radial stretch $\lambda_R(R_0)$, and the incompressibility condition $\lambda_R\lambda_C\lambda_t = 1$, follows that $\lambda_t(R_0) = 1/(\lambda_C\lambda_R)$. The lateral strain energy is then

$$E_{\text{lateral}} = N_{\text{cells}}\frac{1}{6}E_y t_0 \int_{R_{i,0}}^{R_{i,o}} C_0(R_0) W_{\text{lat}}(\lambda_C(R_0), \lambda_R(R_0))\, dR_0 \tag{C19}$$

where $W_{\text{lat}}(\lambda_C, \lambda_R) = \lambda_C^2 + \lambda_R^2 + (\lambda_C\lambda_R)^{-2} - 3$ is the neo-Hookean strain energy density function. This expression can be evaluated numerically (see Fig. C1), but cannot be reduced to a closed form because of the implicit non-linear dependence of the stretches on $R_0$. To obtain a tractable analytical approximation, we assume a uniform radial stretch $\lambda_H$ and circumferential stretch $\lambda_C$ across the shell thickness. This allows us to pull $W_{\text{lat}}$ outside the integral. Defining the average reference circumference,

$$\bar{C}_0 = \frac{1}{H_0}\int_{R_{i,0}}^{R_{i,o}} C_0(R_0)\, dR_0 = \alpha_n\sqrt{\frac{4\pi}{N_{\text{cells}}}}(R_{i,o} + R_{i,0}) \tag{C20}$$

we obtain the approximate energy

$$E_{\text{lateral}} \approx N_{\text{cells}}\frac{1}{6}E_y t_0 \bar{C}_0 H_0 \left[\lambda_C^2 + \lambda_H^2 + (\lambda_C\lambda_H)^{-2} - 3\right] \tag{C21}$$

***Stretch functions***

We can express all stretches in terms of lumen strain $\epsilon = (R_i - R_{i,0})/R_{i,0}$. The inner (apical) stretch is

$$\lambda_i = 1 + \epsilon \ . \tag{C22}$$

The outer (basal) stretch follows from incompressibility of the shell $(R_o^3 - R_i^3)/(R_{o,0}^3 - R_{i,0}^3) = 1$ and the thickness ratio $\beta = (R_{o,0} - R_{i,0})/R_{o,0}$,

$$\lambda_o = \left[1 + (1-\beta)^3(\lambda_i^3 - 1)\right]^{1/3} \ . \tag{C23}$$

The radial stretch is defined by

$$\lambda_H = \frac{H}{H_0} = \frac{R_o - R_i}{R_{o,0} - R_{i,0}} = \frac{\lambda_o}{\beta} - \frac{(1-\beta)\lambda_i}{\beta} \tag{C24}$$

Finally, imposing incompressibility at the single-cell level, with reference volume $V_{\text{cell},0} = d^3 = A_{tan,0}H_0 = A_{\text{tan}}H$, with $A_{\text{tan}}$ a representative tangential (circumferential) area, and noting that $A_{\text{tan}} \sim C^2$, gives

$$\lambda_C = \sqrt{\frac{A_{\text{tan}}}{A_{\text{tan},0}}} = \sqrt{\frac{H_0}{H}} = \lambda_H^{-1/2} \ . \tag{C25}$$

***Pressure function (averaged lateral term)***

Defining pressure as $P = dE_{\text{tot}}/dV_i$, with $V_i = \frac{4}{3}\pi(R_{i,0}(1+\epsilon))^3$. We can get the pressure versus strain as

$$\begin{aligned} P(\epsilon) = \frac{E_y t_0}{3R_{i,0}}\Big[&(2\lambda_i^{-1} - 2\lambda_i^{-7}) + (1-\beta)(2\lambda_o^{-1} - 2\lambda_o^{-7}) \\ &+ L(\beta, R_{i,0})(\lambda_H - \lambda_H^{-2})\left(\frac{1}{\lambda_o^2} - \frac{1}{\lambda_i^2(1-\beta)^2}\right)\Big] \end{aligned} \tag{C26}$$

$$L(\beta, R_{i,0}, d) = \alpha_6(2-\beta)\sqrt{\frac{\beta(3 - 3\beta + \beta^2)}{3(1-\beta)}}\left(\frac{R_{i,0}}{d}\right)^{3/2} \tag{C27}$$

with linear response

$$P_{\text{linear}} = \frac{E_y t_0}{3R_{i,0}} \left[ 12 + 12(1-\beta)^4 + \frac{3\beta(2-\beta)^2}{1-\beta} L \right] \epsilon \tag{C28}$$

In the linear and intermediate regime this function matches numerical evaluation of the full model. At higher strains the theory slightly underestimates the pressure (Fig. C1), due to the non-linearity of $R(R_0)$.

#### C.2 Cellular thin-sheet model with a soft lateral mode

When the lateral faces give way to soft modes, the epithelium can undergo a global change in thickness $H$ but the local radial compression in the lateral sheet becomes negligible. As a consequence the incompressibility condition becomes $\lambda_C \lambda_t = 1$, and $\lambda_t(R_0) = 1/\lambda_C(R_0)$. The full expression has the same shape as Eq. C19, but now $W_{\text{lat}}(\lambda_C(R_0)) = \lambda_C^2 + \lambda_C^{-2} - 2$ (see Fig. C1). For the approximation $\lambda_C$ still scales with global thinning via $\lambda_C = \lambda_H^{-1/2}$ and thus we find,

$$E_{\text{lateral,soft}} \approx N_{\text{cells}} \frac{1}{6} E_y t_0 \bar{C}_0 H_0 \left( \lambda_C^2 + \lambda_C^{-2} - 2 \right) \tag{C29}$$

Introducing the soft mode leads to a minor modification of the pressure equation (see Fig. C1).

$$\begin{aligned} P_{\text{soft}}(\epsilon) = \frac{E_y t_0}{3R_{i,0}} \Big[ &(2\lambda_i^{-1} - 2\lambda_i^{-7}) + (1-\beta)(2\lambda_o^{-1} - 2\lambda_o^{-7}) \\ &+ \frac{1}{2} L(\beta, R_{i,0})(1 - \lambda_H^{-2}) \left( \frac{1}{\lambda_o^2} - \frac{1}{\lambda_i^2 (1-\beta)^2} \right) \Big] \end{aligned} \tag{C30}$$

with linear response

$$P_{\text{soft,linear}} = \frac{E_y t_0}{3R_{i,0}} \left[ 12 + 12(1-\beta)^4 + \frac{\beta(2-\beta)^2}{1-\beta} L \right] \epsilon \tag{C31}$$

Comparison of the linearized soft-mode model and full model expressions shows a shift in the relative contributions of the different faces to the resistance to inflation. For lateral compression, the contributions are Apical 41%, Basal 3%, and Lateral 56%. For the lateral soft mode, they are Apical 66%, Basal 4%, and Lateral 30%. These estimates assume uniform cortical thickness and modulus across all faces. In absolute terms, the lateral contribution in the soft mode is about one-third of that in the compression mode.

#### C.3 Comparison with experiment and assumptions

The analytical expressions for the cellular models rely on several assumptions:

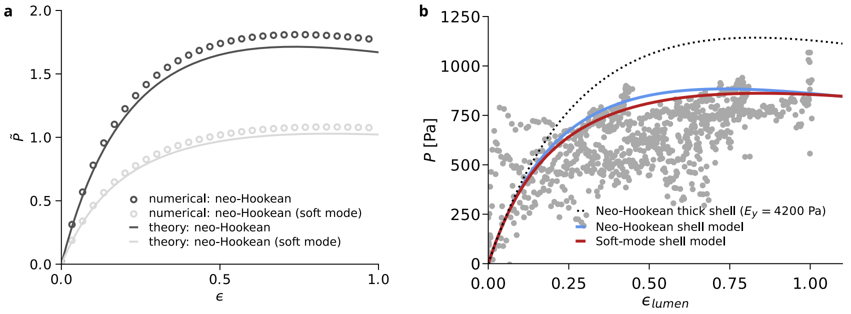


**Fig. C1** [a] Normalized pressure ($\tilde{P} = \frac{P}{E_y}\frac{R_{i,0}}{t_0}$) versus luminal strain for inflation of a 3D shell. Theoretical curves for the neo-Hookean model and the soft mode are based on Eq. C26 and Eq. C30, respectively. The numerical curves are based on evaluation of $dE/dV$ of the total energy Eq. using the full lateral energy expression Eq. C19. [b] Prediction of non-linear response from thick-shell and cellular thin-sheet models based on linear fit to experimental data (grey dots).

- **Material model:** Elasticity is dominated by the actomyosin cortex (negligible effect of nucleus and intermediate filament networks). The cortex is taken as incompressible, with spatially uniform stiffness and thickness across all faces, and assumed to respond linearly (neo-Hookean), i.e. negligible strain stiffening.
- **Geometry and kinematics:** Each face is treated as a thin sheet with homogeneous stretch across its thickness (no bending energy from curvature). Circumferential stretch of the lateral faces is assumed homogeneous along the radial direction. The epithelium is taken to be spherically symmetric and isotropic, with fixed topology (no cell division or rearrangement).
- **Neglected effects:** Elastic contributions from cell–cell adhesion are ignored, as are viscoelastic and poroelastic contributions to the measured pressure.
- **Reference state:** No active prestress from actomyosin contractility and no passive prestretch are included; the reference configuration is stress-free.

The epithelial mechanics explored in our experiments and models represents a limiting case. Nevertheless, this limit follows naturally from more general formulations of epithelial mechanics that include viscous and active contributions. [6, 7] In particular, recent frameworks coupling hyperelasticity and active tension [7], reduces to a purely elastic description if the deformation is fast compared to viscous and active timescales. Our experiments therefore provide a direct probe of the hyperelastic response of the actin cortex, which may serve as a useful reference for mesoscale models of cytoskeletal mechanics and thus help quantify the impact of structural defects on tissue-scale mechanics.

This connection also enables direct comparison between our fitted parameters and values reported in literature. Ouzeri et al. [7], for instance, fit uniaxial stretching data of MDCK epithelia [6] by setting the 2D shear modulus $\mu_{2D}$ equal to the steady-state active tension, 0.975 mN/m. For our hiPSC data we obtain $\mu_{2D} = 2.75$ mN/m from the full model and $\mu_{2D} = 4.47$ mN/m when soft modes are considered. These values are of the same order of magnitude. The higher modulus in hiPSCs may reflect structural differences; in particular, MDCK cells lack prominent actin belts, which could account for part of the discrepancy.

### D discrete model for 2D cortinoids

#### D.1 Numerical approach

We construct a 2D model of a ring of cells specified by the characteristic cell size $d$. Each cell consists of vertices connected by edges, and displacement of these vertices incurs a harmonic energy penalty from four contributions: resistance to area change ($k_A$), edge stretching ($k_s$), whole-cell tilt ($k_t$), and lateral-edge bending ($k_b$). To allow for bending we divide the lateral edge in 20 equally sized segments.

The half width of each cell in radians is $\Delta\alpha = \pi/N_{\text{cells}}$. We introduce geometrical disorder by independently modulating the tangential coordinate of the inner and outer nodes as $\Delta\alpha \to \Delta\alpha\,[1 + U(-\delta, \delta)]$, with $U(-\delta, \delta)$ a uniform random variable, which produces variability in tilt and cell size.

To model the inflation, we constrain the nodes on the outer edge of this ring to a growing circle. Furthermore we fully fix the radial angle of one of the nodes on the outer-ring to suppress zero energy global rotation mode. We take a quasistatic approach, where we determine the minimum energy configuration as a function of the radius of this circle. The energy is defined as follows:

$$\mathcal{E} = \mathcal{E}_A + \mathcal{E}_S + \mathcal{E}_T + \mathcal{E}_B \tag{D32}$$

We define $\mathcal{E}_A$ as

$$\mathcal{E}_A = \frac{1}{2}\frac{k_A}{A_0}\sum_{cells}(A - A_0)^2 \quad . \tag{D33}$$

We define $\mathcal{E}_S$ as

$$\mathcal{E}_S = \frac{k_S}{2}\sum_{cells}\left[\frac{(l - l_{a,0})^2}{l_{a,0}} + \frac{(l - l_{b,0})^2}{l_{b,0}} + \sum_{lateral}\frac{(l - l_{l,0})^2}{l_{l,0}}\right] \quad . \tag{D34}$$

with distinct contributions for the apical, basal, and lateral edges.

We define $\mathcal{E}_T$ as

$$\mathcal{E}_T = \sum_{cells}\sum_{\langle ijk\rangle_{corner}}\frac{k_t}{l_{0,ij} + l_{0,jk}}(\theta - \theta_{0,ijk})^2 \quad . \tag{D35}$$

We define $\mathcal{E}_B$ as

$$\mathcal{E}_B = \sum_{cells}\sum_{\langle ijk\rangle_{lateral}}\frac{k_b}{l_{0,ij} + l_{0,jk}}(\theta - \theta_{0,ijk})^2 \quad . \tag{D36}$$

We find the minimum energy configuration through the minimization of these energies with the Fast Inertial Relaxation Engine (FIRE)[8] through an adapted version of the protocol in JAX-MD[9] implementing the constraints described above as well as a modification towards FIRE 2.0[10]. We use the auto-differentiation functionality of JAX to retrieve the forces from the energies described above. The parameters for FIRE minimization are $dt_{\text{start}} = 0.001$, $dt_{\text{min}} = 0.001$, $dt_{\text{max}} = 0.01$, $N_{\text{min}} = 5$, $f_{\text{inc}} = 1.1$, $f_{\text{dec}} = 0.5$,$\alpha_{\text{start}} = 0.1$, $f_\alpha = 0.99$. In all cases $F_{rms} = 1 \times 10^{-9}$.

We retrieve the pressure from the slope of the energy versus area curves, $P = dE/dA$. We calculate the energy of the system upon deformation by a small strain step below and above the current strain. To reduce the number of free parameters, we fix $k_s = 1$ and $d = 1$, define $\tilde{k}_t = k_t/(k_s d^2)$, $\tilde{k}_b = k_b/(k_s d^2)$, $\tilde{k}_A = k_A d/k_s$, and enforce effective incompressibility by $\tilde{k}_A \gg 1$. Extende Fig. 3 displays the range of mechanical responses that are produced by the model.

Strain-steps are imposed through areal strain $\epsilon_A$ of the entire cortinoid. For mechanical response curves we use 150 linearly spaced steps from $\epsilon_A = 0$ to 0.75. For onset identification we use 150 logarithmically spaced steps from $\epsilon_A = 1 \times 10^-3$ to 0.75. Note that we always report luminal strain $\epsilon = \sqrt{1 + \epsilon_A/(1-\beta)^2} - 1$, to facilitate comparison across experiment and models.

#### D.2 Mechanics: Stretch based limits

Following a similar procedure as for the 3D thin-faces model, we can derive analytical expressions for the mechanical limits of the 2D model.

$$E_{\text{tot}} = E_{\text{apical}} + E_{\text{basal}} + E_{\text{lateral}} \tag{D37}$$

We define strains on the basal edges ($\epsilon_o$) and on the lateral edges ($\epsilon_H$) in terms of the lumen strain $\epsilon = (R_i - R_{i,0})/R_{i,0}$ and the geometric parameter $\beta = (R_{o,0} - R_{i,0})/R_{o,0}$. Imposing area incompressibility of the 2D cells, $\pi R_o^2 - \pi R_i^2 = \pi R_{o,0}^2 - \pi R_{i,0}^2$, relates the deformed outer radius $R_o$ to the lumen strain $\epsilon$. The basal strain then becomes

$$\epsilon_o = \frac{R_o - R_{o,0}}{R_{o,0}} = \sqrt{1 + (1-\beta)^2(\epsilon^2 + 2\epsilon)} - 1 \tag{D38}$$

the lateral strain follows from the relative displacement of the inner and outer boundaries,

$$\epsilon_H = \frac{H - H_0}{H_0} = \frac{(R_o - R_i) - (R_{o,0} - R_{i,0})}{R_{o,0} - R_{i,0}} = \frac{\epsilon_o - (1-\beta)\epsilon}{\beta} \tag{D39}$$

For the apical and basal faces we can write the energy

$$E_{\text{apical}} = \frac{k_s}{2L_0}(L - L_0)^2 = \pi R_{i,0} k_s \epsilon^2 \tag{D40}$$

$$E_{\text{basal}} = \frac{k_s}{2L_0}(L - L_0)^2 = \pi R_{o,0} k_s \epsilon_o^2 \tag{D41}$$

$$E_{\text{lateral}} = N_{\text{cells}} k_s H_0 \epsilon_H^2 \tag{D42}$$

We can define the pressure as $P = dE_{\text{tot}}/dA_i$, with $A_i = \pi R_{i,0}^2(1+\epsilon)^2$, leading to

$$P = \frac{k_s}{R_{i,0}}\left[\frac{\epsilon}{1+\epsilon} + (1-\beta)\frac{\epsilon_o}{1+\epsilon_o} + \frac{R_{i,0}^2}{d^2}\frac{\beta(2-\beta)}{(1-\beta)}\epsilon_H\left(\frac{1}{1+\epsilon_o} - \frac{1}{(1-\beta)(1+\epsilon)}\right)\right] \tag{D43}$$

with

$$P_{\text{linear}} = \frac{k_s}{R_{i,0}}\left[1 + (1-\beta)^3 + \frac{R_{i,0}^2}{d^2}\frac{\beta^2(2-\beta)}{(1-\beta)}\right]\epsilon \tag{D44}$$

To get the expressions in the soft mode limit, we just remove the third (lateral) term completely. Extended Fig. 3 reveals that the full model corresponds to the upper limit of the mechanical response that can be obtained through the simulations, while the soft mode limit (excluding the third term between brackets), corresponds to the lower limit.

#### D.3 Geometry: Scaling of bend and tilt

***Minimal model for lateral tilting***

In the absence of the tilt and bend resistance, the system allows zero energy mode of deformation corresponding to parallel displacement of the apical and basal faces. In the idealized flat and symmetric limit($\beta \to 0$, $\delta = 0$), where cells have height and width $d = H_0$ (see Extended Fig. 4, we can derive an analytical expression for the tilt angle that preserves the lateral edge length.

$$\theta(\epsilon) = \arccos\left(\frac{1}{1+\epsilon}\right) \sim \sqrt{2\epsilon}, \quad \epsilon \to 0^+ \tag{D45}$$

To assess the effect of tilt resistance in this minimal setting, we use an energy minimization argument. For a given sheet height $H$, the lateral angle $\theta$ may vary between 0 and $\cos^{-1}(H/H_0)$, with $l_{\text{lat}} = H/\cos(\theta)$ and $H = H_0/(1+\epsilon)$. The total energy including resistance to tilt and lateral edge compression is

$$E_s + E_t = \frac{k_s}{2H_0}\left(\frac{H}{\cos\theta} - H_0\right)^2 + \frac{k_t}{H_0}\theta^2 \tag{D46}$$

Note that to allow a direct comparison with our simulations, we consider that for each lateral edge there are two tilt angle contributions, from the apical and basal side respectively. This system is stationary when $dE/d\theta = 0$, i.e.,

$$\frac{k_s}{H_0}\left(\frac{H}{\cos\theta} - H_0\right)\frac{dl_{\text{lat}}}{d\theta} + 2\frac{k_t}{H_0}\theta = 0 \tag{D47}$$

with $dl_{\rm lat}/d\theta = H\sin\theta/\cos^2\theta$. Rearranging gives

$$\tilde{k}_t \equiv \frac{k_t}{k_s H_0^2} = \frac{1}{2}\frac{\sin\theta}{\theta}\frac{(1+\epsilon)\cos\theta - 1}{(1+\epsilon)^2\cos^3\theta} \tag{D48}$$

since $H_0 = d$, the left-hand side scales with $\tilde{k}_t$. This relation predicts a pitchfork bifurcation for $\tilde{k}_t > 0$, comprising a trivial untilted branch ($\theta = 0$) and a pair of symmetry-related tilted branches ($\theta = \pm\theta(\epsilon)$) that emerge at a critical strain $\epsilon_c$. To characterize the behaviour near this onset, we expand Eq. D48 around $\epsilon_c$ and $\theta = 0$. Writing $\epsilon = \epsilon_c + \Delta\epsilon$, with $\Delta\epsilon \ll 1$, the right-hand side can be Taylor expanded in $\theta$ around $\theta = 0$ as

$$\frac{1}{2}\frac{\sin\theta}{\theta}\frac{(1+\epsilon)\cos\theta - 1}{(1+\epsilon)^2\cos^3\theta} = \frac{1}{2}\frac{\epsilon}{(1+\epsilon)^2} + \frac{1}{2}A(\epsilon)\theta^2 + O(\theta^4) \tag{D49}$$

with $A(\epsilon) = (5\epsilon - 3)/(6(1+\epsilon)^2)$. This yields non-zero solutions for $\theta$ beyond a critical strain. At the same strain, the trivial branch loses linear stability. Since at onset $\theta = 0$, the critical strain follows from $\tilde{k}_t = \frac{1}{2}\epsilon_c/(1+\epsilon_c)^2$. As the right-hand side attains its maximum at $\epsilon_c = 1$, this equation has solutions only if $\tilde{k}_t \leq 1/8$; for $\tilde{k}_t > 1/8$ no symmetry-breaking tilt occurs. For $\epsilon > \epsilon_c$ the balance becomes

$$2\tilde{k}_t - \frac{\epsilon}{(1+\epsilon)^2} \approx A(\epsilon_c)\theta^2 \tag{D50}$$

The sign of $A(\epsilon_c)$ determines the bifurcation type. For $\epsilon_c < 3/5$ ($A < 0$) the non-trivial solution grows continuously for $\epsilon \to \epsilon_c^+$ (supercritical), whereas for $\epsilon_c > 3/5$ ($A > 0$), finite-tilt states already exist for some $\epsilon < \epsilon_c$ (subcritical). Since $\epsilon_c = 3/5$ corresponds to $\tilde{k}_t = \frac{15}{128}$, the bifurcation is supercritical for $\tilde{k}_t < \frac{15}{128}$, in which case

$$\theta(\epsilon) \sim \sqrt{\alpha(\epsilon - \epsilon_c)}\ , \qquad \epsilon \to \epsilon_c^+, \tag{D51}$$

with $\alpha = 6(\epsilon_c - 1)/((1+\epsilon_c)(5\epsilon_c - 3))$. Equivalently, in terms of $T \equiv \theta^2$

$$T(\epsilon) \sim \alpha(\epsilon - \epsilon_c)\ , \qquad \epsilon \to \epsilon_c^+, \tag{D52}$$

Thus, to summarize, we expect supercritical tilting behaviour for $\tilde{k}_t < 15/128$, subcritical behaviour for $15/128 < \tilde{k}_t < 1/8$, and no tilting for $\tilde{k}_t > 1/8$. Extended Fig. 4 shows $T$ versus $\epsilon$ for this flat sheet model. We find that the numerical solutions based on Eq. D48 coincide with the predicted onset (grey dotted lines). We also observe good agreement with the linear approximation in Eq. D52 over most of the strain range (not shown, as the curves are visually indistinguishable).

***Minimal model for lateral bending***

In the absence of bending resistance, the lateral edge can accommodate parallel displacement of the apical and basal faces by buckling without energetic cost. For pinned–pinned boundaries, the lowest buckling mode is

$$w(z) = a \sin\left(\frac{\pi z}{H}\right), \qquad z \in [0, H]. \tag{D53}$$

with transverse displacement $w$, lateral coordinate $z$ along the deformed edge, and amplitude $a$, giving an arc length

$$l_{\text{lat}} \approx H + \frac{\pi^2}{4H} a^2, \tag{D54}$$

for small slopes $(|a\pi/H|) \ll 1$ and stretching energy

$$E_s = \frac{k_s}{2H_0}\Big(l_{\text{lat}} - H_0\Big)^2 = \frac{k_s}{2H_0}\left(-\frac{H_0\epsilon}{1+\epsilon} + \frac{\pi^2(1+\epsilon)}{4H_0} a^2\right)^2. \tag{D55}$$

Taking the discretized bending term Eq. D36 in the continuum limit ($l \to 0$), the small-slope relation $\Delta\theta \simeq \kappa\,\ell$ gives

$$E_b \;\approx\; \frac{k_b}{2}\int_0^{H_0} \kappa^2\, ds_0, \tag{D56}$$

The integral naturally leads to the dimensionless expression $B = H_0 \int \kappa^2 ds$, which we use to quantify our experiments and simulations, and underlines that $B$ is proportional to the bending energy density. For the sinusoidal mode,

$$E_b = \frac{k_b \pi^4}{4H_0^3} a^2. \tag{D57}$$

Stationarity of $E = E_s + E_b$ with respect to $a$ ($dE/da = 0$), then yields

$$\frac{k_s}{H_0}\left(-\frac{H_0\epsilon}{1+\epsilon} + \frac{\pi^2(1+\epsilon)}{4H_0} a^2\right)\frac{\pi^2(1+\epsilon)}{2H_0} a + \frac{k_b\pi^4}{2H_0^3} a = 0. \tag{D58}$$

As for tilt, this expression predicts a pitchfork bifurcation with a trivial branch ($a = 0$) and a non-trivial branch $a = \pm a(\epsilon)$. To extract the non-trivial branch ($a \neq 0$), we can rearrange this expression as

$$\tilde{k}_b \equiv \frac{k_b}{k_s H_0^2} = \frac{\epsilon}{\pi^2} - \frac{(1+\epsilon)^2}{4}\left(\frac{a}{H_0}\right)^2 \tag{D59}$$

revealing the connection to $\tilde{k}_b$, with $H_0 = d$. This yields non-zero solutions for $a$ beyond a critical strain. At the same strain, the trivial branch loses linear stability. At onset ($a = 0$) this gives following relation between $\tilde{k}_b$ and the critical strain $\epsilon_c$

$$\tilde{k}_b = \frac{\epsilon_c}{\pi^2} \tag{D60}$$

Since the right-hand side scales linearly with $\epsilon_c$ there is no upper bound for the onset of the soft-mode as we observed for tilt. To determine the evolution of $a$ beyond the onset we rewrite Eq. D59 as

$$a^2 = \frac{4H_0^2}{\pi^2(1+\epsilon)^2}\left[\epsilon - \pi^2\tilde{k}_b\right] = \frac{4H_0^2}{\pi^2(1+\epsilon)^2}\left[\epsilon - \epsilon_c\right] \tag{D61}$$

Linearizing as $\epsilon = \epsilon_c + \Delta\epsilon$ with $\Delta\epsilon \ll 1$, we can approximate this expression as

$$a^2 \approx \frac{4H_0^2}{\pi^2}\frac{\epsilon - \epsilon_c}{(1+\epsilon_c)^2}. \tag{D62}$$

so that

$$a(\epsilon) \sim \frac{2H_0}{\pi}\frac{1}{(1+\epsilon_c)}\sqrt{\epsilon - \epsilon_c}, \qquad \epsilon \to \epsilon_c^+. \tag{D63}$$

or in terms of $B$,

$$B(\epsilon) \approx \frac{2\pi^2}{(1+\epsilon_c)^2}(\epsilon - \epsilon_c), \qquad \epsilon \to \epsilon_c^+. \tag{D64}$$

Since the slope is positive $(2\pi^2/(1+\epsilon_c)^2 > 0)$, the buckling amplitude grows continuously from zero once the critical strain is exceeded, i.e. the bifurcation is supercritical for all $\tilde{k}_b$. Extended Fig. 4 shows $B$ versus $\epsilon$ for the flat sheet model. The numerical solutions based on Eq. D59 coincide with the predicted onset (grey dotted lines). We observe good agreement with the linear approximation in Eq. D64 over most of the strain range (not shown, as the curves are visually indistinguishable).

***Interpretation of 2D simulations***

In Extended Fig.4 we compare the 2D flat sheet theory with simulations of a 2D cortinoids. For the fully symmetric case ($\delta = 0$) the simulated response qualitatively matches the theory. Small shifts in the onset strain arise from (i) the non-linear mapping of lateral strain on radial strain in a spherical geometry, ii) the larger cell-height in the simulations ($H_0 = 1.6d$ for $\beta = 0.5$ and $R_{i,0} = 1.6d$). Apparent tilt beyond the flat-sheet limit in simulations is likewise explained by the larger $H_0$. The theory uses $\tilde{k} \propto 1/H_0^2$ whereas the simulations use $\tilde{k} \propto 1/d^2$), shifting thresholds by $(H_0/d)^2$. Introducing disorder in the initial tilt distribution ($\delta = 1$), noticeably smooths the crossover in $B$ and $T$. For tilt, part of the difference stems from the observable $T$ (red), which is influenced by the initial offsets. Plotting $\langle(\theta - \theta_0)^2\rangle$ (grey) removes this bias but significant smoothing of the crossover remains. Disorder does not appreciably affect the location of the onset.

Note that in 3D bend and tilt lead to more complex deformation patterns at the sheet level.
